# A new chemoenzymatic semisynthetic approach provides novel insight into the role of phosphorylation beyond exon1 of Huntingtin and reveals N-terminal fragment length-dependent distinct mechanisms of aggregation

**DOI:** 10.1101/2021.03.24.436743

**Authors:** Rajasekhar Kolla, Pushparathinam Gopinath, Jonathan Ricci, Andreas Reif, Iman Rostami, Hilal A. Lashuel

## Abstract

Huntington’s disease is a neurodegenerative disorder caused by the expansion of a polyglutamine (poly Q) repeat (>36Q) in the N-terminal domain of the huntingtin protein (Htt), which renders the protein or fragments thereof more prone to aggregate and form inclusions. Although several Htt N-terminal fragments of different lengths have been identified within Htt inclusions, most studies on the mechanisms, sequence, and structural determinants of Htt aggregation have focused on the Htt exon1 (Httex1). Herein, we investigated the aggregation properties of mutant N-terminal Htt fragments of various lengths (Htt171, Htt140, and Htt104) in comparison to mutant Httex1. We also present a new chemoenzymatic semisynthetic strategy that enables site-specific phosphorylation of Htt beyond Httex1. These advances yielded novel insights into how PTMs and structured domains beyond Httex1 influence aggregation mechanisms, kinetics, and fibril morphology of longer N-terminal Htt fragments. We demonstrate that phosphorylation at T107 significantly slowed its aggregation, whereases phosphorylation at T107 and S116 accelerated the aggregation of Htt171, underscoring the importance of crosstalk between different PTMs. We demonstrate that mutant Htt171 proteins aggregate via a different mechanism and form oligomers and fibrillar aggregates with morphological properties that are distinct from that of mutant Httex1. These observations suggest that different N-terminal fragments could have distinct mechanisms of aggregation and that a single polyQ-targeting anti-aggregation strategy may not effectively inhibit the aggregation of all N-terminal Htt fragments. Finally, our results underscore the importance of further studies to investigate the aggregation mechanisms of Htt fragments and how the various fragments interact with each other and influence Htt toxicity, pathology formation, and disease progression.

**Table of content:** 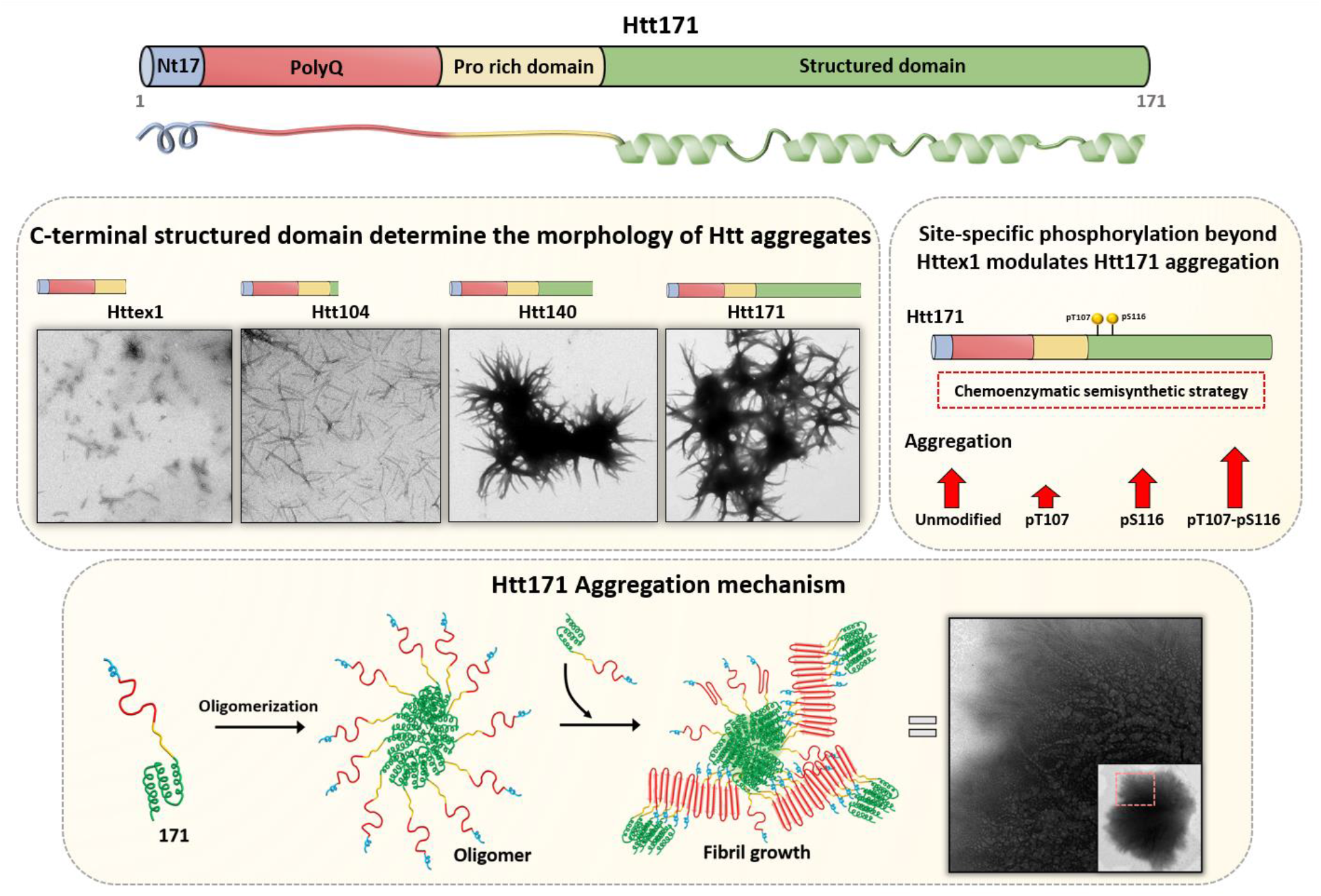

## Introduction

Huntington’s disease (HD) is an inherited progressive neurodegenerative disorder causing movement disabilities, personality changes, and cognitive impairment.^1,2^ At the pathological level, HD is characterized by the presence of proteinaceous inclusions composed of aggregated Huntingtin protein (Htt) proteins with an expanded polyglutamine (poly Q) repeat stretch of >36 glutamine residues, in the affected brain regions.^3,4^ Htt is expressed as a 3144 amino acid (AAs) protein and is subject to several post-translational modifications (PTMs), including phosphorylation, acetylation, polyamidation, sumoylation, ubiquitination, and proteolysis.^5^ These PTMs are distributed throughout the Htt protein sequence and predominantly exist as clusters in the unstructured regions of the protein.^6^ Previous studies on Htt have focused primarily on the PTMs within the first 17 N-terminal residues of the protein because of the identification of several types of PTMs in this region and their proximity to the disease-causing and aggregation-prone poly Q domain.^7^ Several other phosphorylation sites beyond Httex1 (S421, S434, S536 S1181, S1201, S2076, S2653, and S2657) were identified and shown to modulate different aspects of Htt function, cellular properties, degradation and toxicity (Figure 1).^8,9,10,11,5^ Moreover, recently mass spectrometry analyses of HD mouse and human post-mortem brains identified several novel phosphorylation sites in different regions of the Htt sequence.^12^ Two of these phosphorylation sites, S116 and S120, are of special interest because they occur in close proximity to two sites: (1) the putative proteolytic cleavage sites that are thought to contribute to the generation of the highly aggregation-prone protein and toxic Httex1;^13,14^ and (2) two aggregation-prone sequence motifs in exon2/3 (Htt105-116 and Htt128-138).^15^ In addition, blocking phosphorylation at S116 was shown to have a protective effect against the mutant Htt-induced cellular toxicity, suggesting a possible role for phosphorylation at S116 in modulating Htt toxicity.^16^ Interestingly, mimicking or blocking phosphorylation at T107 did not significantly alter Htt586 toxicity. Although these studies suggest a critical role for phosphorylation at S116, they were based on assessing the effect of phosphomimicking mutations, which we have previously shown not to reproduce all aspects of *bona fide* phosphorylation effects.^17,18,19,20^ Recently, expression of the nemo-like kinase (NLK), which phosphorylates Htt at S120, was demonstrated to result in selective lowering of mutant Htt levels in mammalian cells and HD mouse models.^21^ Although NLK phosphorylates Htt at S120, both the specificity and efficiency of this kinase toward Htt remain unknown. Therefore, elucidating the role of phosphorylation at these residues and their relative contributions to modulating the structure and function of Htt and its fragments in healthy and disease states requires the development of methodologies that enable homogeneous and site-specific phosphorylation of Htt at these residues. Our laboratory has developed several semisynthetic and chemoenzymatic strategies for the introduction of single or multiple PTMs in the Nt17 domain of Httex1.^17,22,18,23,24^ Notably, prior to the work described here, there were no methods that made this possible in domains beyond Httex1, for example, in longer N-terminal Htt fragments. Thus, precluding systematic studies to elucidate the role of PTMs, such as T107, S116, and S120, in regulating Htt structure and aggregation and/or potential crosstalk between these phosphorylation sites and neighboring PTMs (such proteolytic cleavage site between C105-A114^13^).

**Figure 1.**
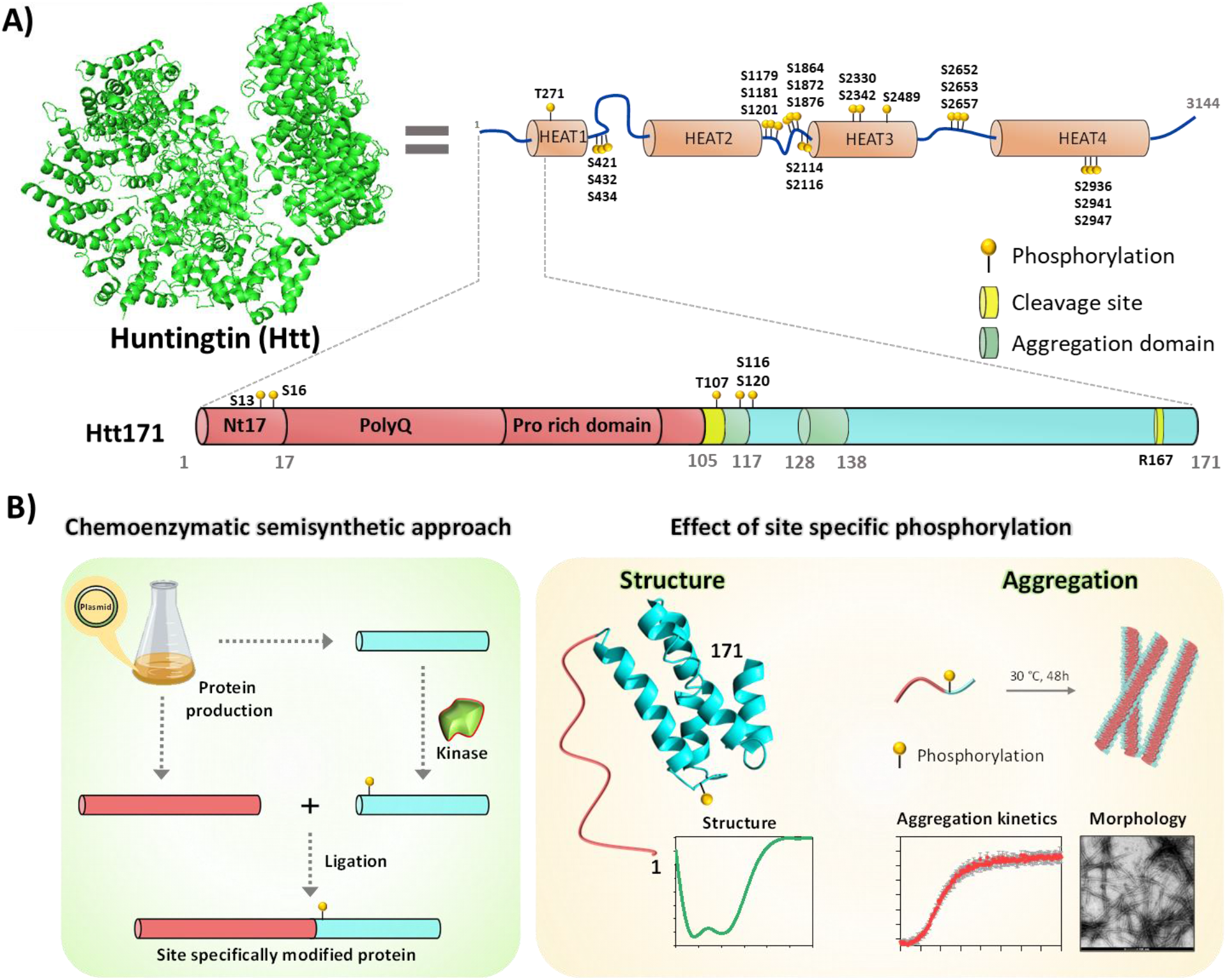
A) Cryo electron microscopy (EM) structure (PDB: 6EZ8)^31^ of huntingtin (Htt) protein and schematic representation of Htt protein with **H**untingtin, **e**longation factor 3, protein phosphatase 2 PP2**A**, and yeast kinase **T**OR1 (HEAT) domains. Htt171 contains an N-terminal Nt17 region, poly Q, proline-rich domain and C-terminal structured domain. The proteolytic cleavage sites, clusters of phosphorylation sites identified in Huntingdon’s disease (HD) mice and human post-mortem brains, phosphorylation at T107 and aggregation domain are highlighted. B) A schematic depiction of the chemoenzymatic semisynthetic approach used in this study to generate site specifically phosphorylated Htt171 proteins and methods to assess the effect of the phosphorylation sites on the structure and aggregation of Htt171.

Herein, we describe the development of a novel chemoenzymatic strategy that addresses the technical challenge mentioned above and enables for the first time, the introduction of single or multiple phosphorylation sites within exon3, present in one of the most commonly studied longer Htt N-terminal fragments, Htt171.^25,26^ The Htt171 fragment was chosen for several reasons: (1) overexpression of mutant Htt171 results in inclusion formation in cellular and animal models of HD, some of which recapitulate a number of behavioural and pathological abnormalities observed in HD pathology^26,27,28,29^ 2) identification of two proteolytic sites, Cp1 (Htt91-114) and Cp2 (R167) in the Htt171 fragment^14^, makes it an ideal model for future studies to explore the role of PTMs in regulating Htt cleavage and the generation of the highly toxic and aggregation-prone Httex1; (3) unlike Httex1, Htt171 has a structured domain (Htt96-171), thus providing a unique opportunity to investigate for the first time how structured domains outside exon1 influence the aggregation mechanisms and properties of longer mHtt fragments; and (4) Htt171 is one of the most commonly studied N-terminal fragments besides Httex1 and Htt586,^3,30^ which enables us to assess our findings in the context of existing data on the aggregation properties of mutant Htt171 in different cellular and animal models of HD. Semi-synthesis of the Htt171 protein and its phosphorylated variants was facilitated by *in vitro* kinome screening studies that led to the identification of novel kinases that efficiently and quantitatively phosphorylate Htt at T107, S116, or both residues. Thus, facilitating the generation of Htt171 proteins that are homogeneously and site-specifically phosphorylated at T107, S116 or both residues. Unfortunately, the lack of specificity of NLK and failure to identify a kinase that site-specifically and efficiently phosphorylates Htt at S120 precluded efforts to produce Htt171-pS120.

Although the aggregation properties of Htt171 protein have been widely studied in several cellular and animal models,^26,27,28,29^ no studies in the literature on the *in vitro* biophysical characterization of mutant Htt171 or other N-terminal fragments than Httex1 have been done. For the first time, we present a comparative analysis of the aggregation properties of Httex1, Htt171, and two other N-terminal fragments in which either full or part of the structured helical domain corresponding to residues 104-171 (Htt104 and Htt140, respectively) was removed. Collectively, our findings provide novel insight into how PTMs and structured domains outside exon1 could significantly modify the kinetics and aggregation pathway of longer N-terminal Htt fragments in addition to the final morphology of mutant Htt aggregates *in vitro*. Our data revealed that Htt171 aggregation occurs *via* distinct mechanisms where initial oligomerization events are driven by structured domains outside Httex1. Collectively, our findings suggest that a single polyQ-targeting anti-aggregation strategy may not effectively inhibit the aggregation of different N-terminal Htt fragments and that multiple N-terminal Htt substrates should be used in future screenings to identify effective Htt aggregation inhibitors. The new semi-synthetic strategies, protein production methodologies, and novel kinases described in this study pave the way for advancing our understanding of the Htt PTM code beyond Nt17. These advances should also facilitate investigation of the role of phosphorylation in regulating proteolytic cleavage of the N-terminal fragment that could lead to the generation of the pathogenic and highly aggregation-prone Httex1 and offer new, exciting opportunities for targeting PTMs for the treatment of HD.

## Results

### Design and semi-synthesis of Htt171-23Q/43Q proteins

The strategy for the semi-synthesis of the Htt171 protein was based on a two-fragment approach (Figure 2). To preserve the native sequence of Htt171 protein, the native cysteine residue (Cys105) was selected as the ligation site. The two protein fragments were recombinantly expressed comprising residues 2−104 with a C-terminal thioester and 105–171 with N-terminal cysteine, respectively (Figures 2B and S1). The recombinant Htt2-104-Mxe intein was expressed in *Escherichia coli* wherein the intein serves as a surrogate for the generation of the C-terminal thioester. The Htt2-104 was expressed in fusion with Mxe-Intein, which contained a C-terminal hexahistidine (His6) purification tag to facilitate affinity-based removal of Htt2-104-23Q/43Q-Mxe-His6 from the bacterial lysate. The fusion protein (**1**) obtained after the affinity purification was then cleaved using 0.1 M Mesna followed by purification of the thioester by reverse phase-high performance liquid phase chromatography (RP-HPLC) to obtain 3.36 mg and 0.96 mg per liter of culture for Htt2-104-23Q-Mesna (**3a**) and Htt2-104-43Q-Mesna (**3b**), respectively (Figure S1). The lower yield of Htt2-104-43Q was attributed to its high aggregation propensity. The expression and purification of Htt2-104-43Q-Mesna presented a series of challenges, which required extensive optimization of several parameters of the protein expression and purification conditions (supporting information) in order to address these challenges. In addition, during the cleavage step, the hydrolyzed Htt2-104-23Q/43Q-Mesna was observed as a byproduct of purification and could not be isolated separately when using HPLC. Since this Htt2-104-23Q/43Q byproduct was non-reactive, it did not interfere with the ligation reaction and was easy to separate at the end of the ligation reactions. Recombinant Htt105-171 was produced through a small ubiquitin-like modifier (SUMO)-fusion strategy in which the fusion protein His6-SUMO-Htt105-171 (**2**) was expressed and purified via affinity chromatography (Figure S2). The His6-SUMO was cleaved by Ubl-specific protease 1 (Ulp 1)^32,33^ followed by RP-HPLC purification to obtain Htt105-171 (**4**) with a very good yield (6 mg/L of culture). After obtaining these two fragments, a native chemical ligation of **3a** or **3b** with fragment **4** was performed under denaturing conditions. Before the ligation, **3a** or **3b** was treated with trifluoroacetic acid (TFA) to dissolve any preformed aggregates, a process that significantly improved the yield of ligation. The **3a** or **3b** fragment (1.2 equiv) was reacted with **4** (1 equiv), and the reaction was monitored by electrospray ionization-mass spectrometry and ultra-performance liquid chromatography (ESI-MS and UPLC, respectively) as shown in Figure 2C. The native chemical ligation (NCL) proceeded readily, and the ligation was completed in 4 h. The resulting Htt171-23Q (**5a**) or Htt171-43Q (**6a**) protein was purified via RP-HPLC, and its purity was analyzed and confirmed by ESI-MS, UPLC, and sodium dodecyl sulfate polyacrylamide gel electrophoresis (SDS-PAGE) as shown in Figure 2C.

**Figure 2.**
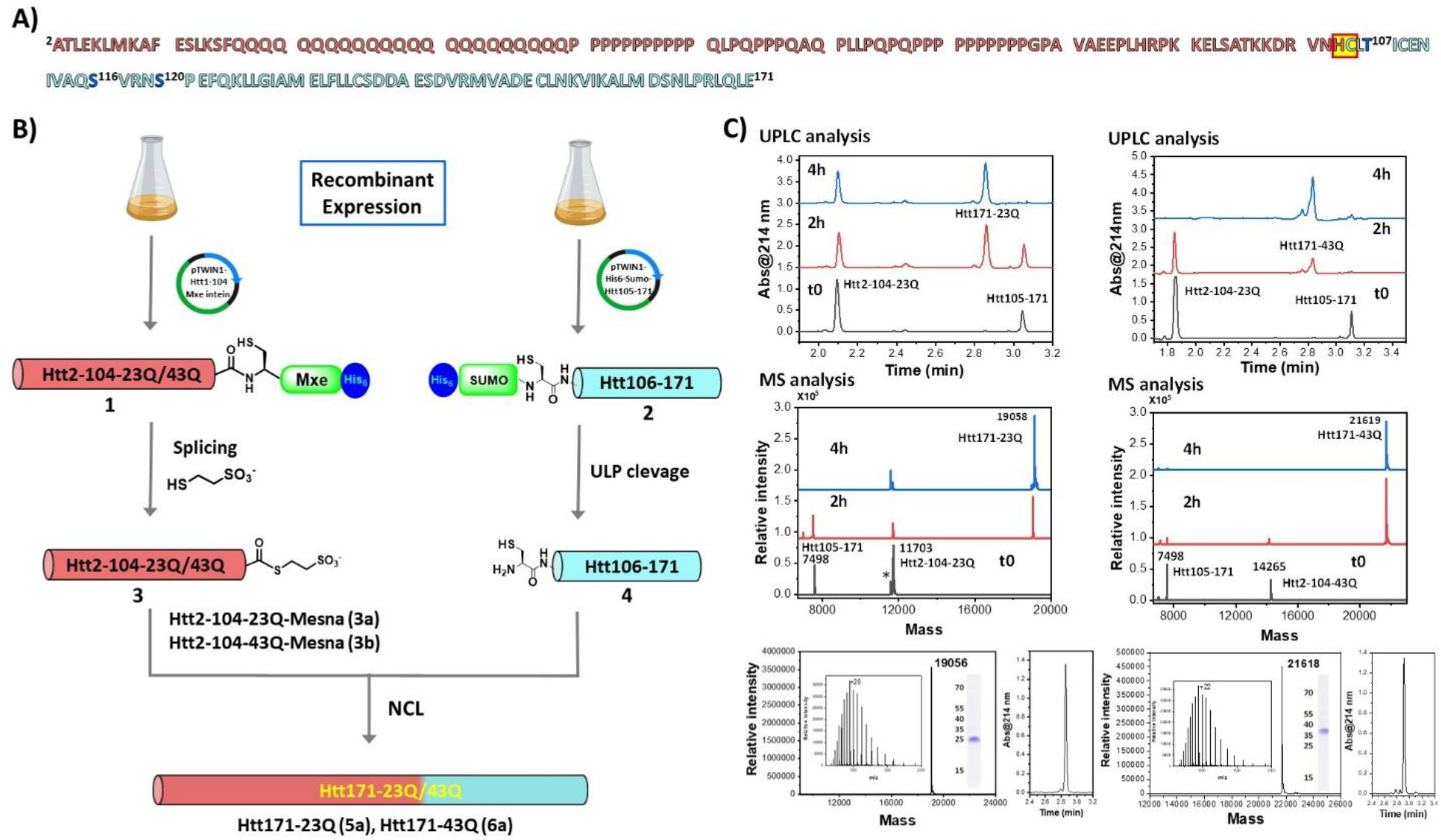
Strategy for the semi-synthesis of Htt171-23Q/43Q. A) Htt171-23Q amino acid sequence with the two fragments shown in red and blue, and the ligation site highlighted by the yellow box. B) Schematic representation for the semi-synthesis of Htt171-23Q/43Q by two fragment approach. C) Ultra-pure liquid chromatography (UPLC) traces and electrospray ionization-mass spectrometry (ESI-MS) analysis of the ligation reaction between the Htt2-104-23Q (left) or Htt2-104-43Q (right) and Htt105-171. Characterization of the purified phosphorylated Htt171-23Q (left) and Htt171-43Q (right) fragments by UPLC, ESI-MS, and sodium dodecyl sulfate polyacrylamide gel electrophoresis (SDS PAGE).

To verify that the semisynthetic Htt171-23Q/43Q proteins fold correctly and adopt their expected conformation, we assessed and compared their secondary structure to the corresponding recombinant Htt171-23Q/43Q proteins (produced in *E. coli*) by circular dichroism (CD) (Figures S3 and S5). Both semisynthetic and recombinant Htt171-23Q/43Q proteins exhibited identical CD spectra and mobility on SDS-PAGE.

### Semi-synthesis of site-specifically phosphorylated Htt171-23Q/43Q proteins

To access the Htt171 proteins with authentic phosphorylation at T107, S116, or S120, site-specific phosphorylated Htt105-171 fragments had to be generated (Figure 3A). To introduce authentic phosphorylation and obtain the phosphorylated 105-171 fragments, we designed two possible approaches: (1) solid-phase peptide synthesis (SPPS) and 2) *in vitro* phosphorylation using kinases that specifically and efficiently phosphorylate Htt at these sites. For the SPPS approach, we designed a two-fragment strategy to access the phosphorylated Htt105-171 fragments (Figure S6). The Htt105-137 (with site-specific phosphorylation at T107, S116, or S120) could be synthesized via SPPS, whereas the Htt138-171 fragment could be generated using SPPS or bacterial expression. Initially, we attempted to synthesize Htt105-137-pS116 and Thz-Htt106-137-Nbz-pT07 using fluorenylmethoxycarbonyl (Fmoc) chemistry. We encountered several difficulties when using SPPS to produce Htt105-137-pS116 (Supporting Information, Figure S7 and S8). To overcome these challenges, several modifications in the synthesis were introduced: (1) changing the solvent system, (2) double coupling, 3) introduction of a pseudo-proline dipeptide to increase solvation of the peptide and improve the coupling efficiency during the synthesis, and 4) using a microwave peptide synthesizer. Although the chemical synthesis of the Thz-Htt106-137-Nbz-pS116 and Thz-Htt106-137-Nbz-pT07 peptides were completed, the ESI-MS analysis showed the presence of several impurities (Figures S7 and S8). Moreover, the crude products obtained from both syntheses exhibited very low solubility, which hindered peptide purification. The insoluble nature of the synthesized Htt105-138 fragment could be explained by the presence of two hydrophobic and highly aggregation-prone domains (Htt108-116 and Htt128-138) [15]. Due to the low yield and solubility of the crude peptide, we had to resort to a different strategy to produce the site-specifically phosphorylated Htt105–171 fragments. This strategy was based on identification of kinases that efficiently phosphorylate Htt at T107, S116, or S120 and then performing an *in vitro* phosphorylation reaction using these kinases to produce Htt105-171 fragments that are site-specifically phosphorylated at each of these residues.

**Figure 3.**
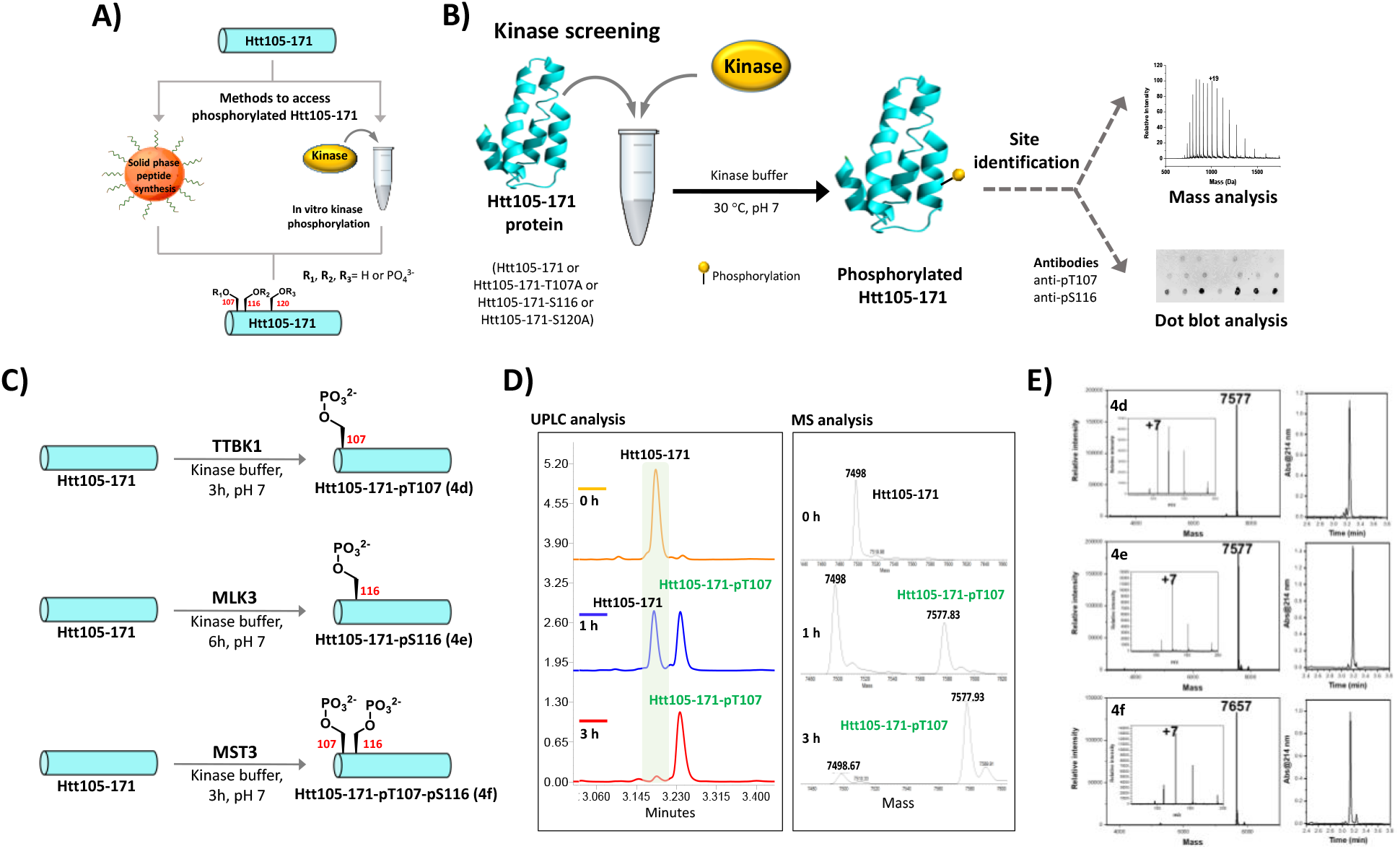
Strategy for accessing phosphorylated Htt105-171 fragments and *in-vitro* kinase reactions A) Strategy for accessing the phosphorylated Htt105-71 fragments through SPPS or *in-vitro* kinase reaction. B) Schematic representation for kinase screening. Protein concentration of 0.5 ug/ul in 60 uL kinase reaction buffer and protein to kinase ratio of 30:1 was maintained C) Schematic representation of kinase reaction for securing Htt105-171-pT107 (**4d**), Htt105-171-pS116 (**4e**) and Htt105-171-pT107-pS116 (**4f**). D) The kinase reaction for **4d** monitored over time by UPLC (left) and ESI-MS (right). E) Characterization of the purified phosphorylated Htt105-171 fragments by analytical ESI-MS and UPLC.

### Kinase screening for identifying site-specifically phosphorylating kinases

The kinases that phosphorylate Htt at T107, S116, or S120 were previously unknown. Therefore, to identify the kinases capable of introducing site-specific phosphorylation at these sites, we performed *in-vitro* kinase screening using a library of 88 kinases (Figure 3B) and Htt105-171 as a substrate. To monitor the extent and site of phosphorylation, the reactions were monitored by ESI-MS and a dot blot analysis using antibodies developed against pT107 and pS116 sites (Figure S9). Unfortunately, at present, no validated antibodies against pS120 exist. To further validate the site of phosphorylation and specificity of the kinases, we generated three recombinant Htt105-171 fragments in which each of these residues was mutated to alanine, Htt105-171-T107A (**4d**), Htt105-171-S116A (**4e**), and Htt105-171-S120A (**4f**) as shown in Figure S4.

The screening was performed concurrently using all the four fragments (**4d–f**), Initially, 20 out of 88 kinases were identified and showed varying degrees of phosphorylation at our desired sites. A careful analysis of the MS data from all the four proteins clearly showed that a handful of kinases phosphorylated either one or two of the desired phosphorylation sites. Among these kinases, the mixed-lineage protein kinase 3 (MLK3) and Tau-tubulin kinase 1 (TTBK1) were shown to selectively and efficiently phosphorylate Htt105-171 at S116 and T107, respectively (Figure S10). The mammalian STE20-like kinase 3 (MST3) selectively and efficiently phosphorylated Htt105-171 at both T107 and S116. Unfortunately, we did not identify any kinases that could selectively phosphorylate at S120. In a recent report, Duan *et al*. reported that NLK phosphorylates Htt at S120 [21]. To determine if NLK could be used to generate the pS120 Htt105-171 fragment, we performed *in vitro* phosphorylation of Htt105-171 proteins (**4d–f**) using recombinant NLK. The MS analysis (Figure S11) showed that NLK phosphorylates Htt105-171 at both S120 and S116. These results indicate that NLK is not site-selective and thus could not be used to generate the Htt105-171-pS120 fragment.

### Generating site specifically phosphorylated Htt105-171 fragments

To determine if the identified kinases could be used to produce good quantities of site-specifically phosphorylated Htt105-171 fragments for use in the ligation reaction for producing the Htt1-171 phosphorylated proteins, we carried out mg scale (Htt105-171) *in vitro* phosphorylation reactions using TTBK1, MLK3, or MST3. The reactions were monitored by ESI-MS and UPLC (Figure 3C and 3D). TTBK1 efficiently phosphorylated Htt105-171at T107 and the reaction was facile (3 h) without any major impurities (Figure 3D). In the case of MLK3, the reaction was complete in 6 h. A small peak corresponding to the di-phosphorylated protein could be detected at later time points but was separated during RP-HPLC purification (Figure S12). MST3 phosphorylated Htt105-171 efficiently at T107 and S116, and the reaction was complete in 3 h (Figure S13). At the initial time points in the reaction, a mixture of mono- and di-phosphorylated forms of Htt105-171 could be observed; however, later in the reaction, the mono-phosphorylated form disappeared, and only the di-phosphorylated Htt105-171 was observed. The resulting proteins, Htt105-171-pT107 (**4a**), Htt105-171-pS116 (**4b**), and Htt105-171-pT107-pS116 (**4c**) were purified by RP-HPLC after which their purities were analyzed and confirmed by ESI-MS and UPLC (Figure 3E).

### Semi-synthesis of site-specifically phosphorylated Htt171-23Q/43Q proteins

After obtaining all of the phosphorylated Htt105-171 fragments, we then focused on assembling the fragments to prepare the desired phosphorylated Htt171 proteins (Figure 4A). The phosphorylated Htt105-171 fragments (**4a–c**) were directly utilized for ligation with Htt2-104-Mesna. TFA-treated **3a** (1.2 equiv) or **3b** (2 equiv) were dissolved in the ligation buffer followed by addition of fragment **4a, 4b**, or **4c** (1 equiv of each). Htt2-104-43Q-Mesna was used in excess (2 equiv) to increase the reaction rate and avoid longer reaction times since the protein is highly prone to aggregation. Similar to the semi-synthesis of the unmodified Htt171 proteins, the ligation went smoothly, and the reaction was completed in 3 to 4 h. The resulting Htt2-171-23Q-pT107 (**5b**), Htt2-171-23Q-pS116 (**5c**), Htt2-171-23Q-pT107-pS116 (**5d**), Htt2-171-43Q-pT107 (**6b**), Htt2-171-43Q-pS116 (**6c**), and Htt2-171-43Q-pT107-pS116 (**6d**) proteins were purified immediately via RP-HPLC, and their purity was confirmed using UPLC, ESI-MS, and Western blotting (Figure 4B). The site of phosphorylation in the phosphorylated Htt171 proteins was validated by dot blot analysis utilizing anti-pT107 and -pS116 antibodies (Figure S14).

**Figure 4.**
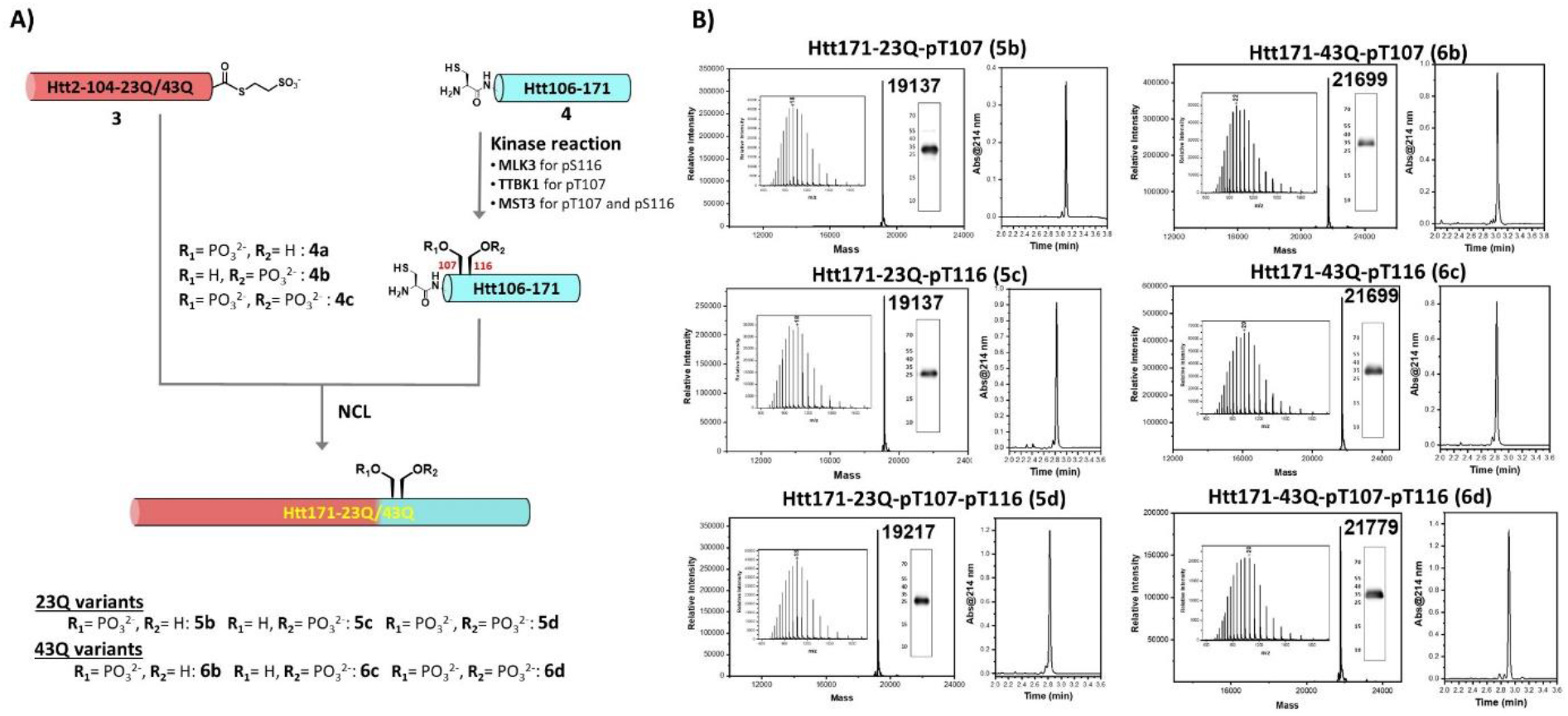
Strategy for the semi-synthesis of phosphorylated Htt171-23Q/43Q proteins. A) Schematic representation for the semi-synthesis of Htt171-23Q/43Q-pT107, Htt171-23Q/43Q-pS116 and Htt171-23Q/43Q-pT107-pS116 by two fragment approach. B) Characterization of the purified Htt171 proteins by analytical ESI-MS, UPLC, and Western blot.

### Htt171-43Q exhibits slower aggregation kinetics in comparison to Httex1-43Q

With all of the phosphorylated and unmodified 171 fragments in hand, we first evaluated the aggregation kinetics (Thioflavin [ThS] assay) and secondary structure (CD analysis) of Htt171-43Q in comparison to Httex1-43Q (Figure 5). Httex1-43Q showed faster aggregation kinetics without a lag phase, a result that also correlated with a shift in the CD spectrum observed after 24h h and indicated transition in secondary structure from a disordered to a β-sheet-rich conformation .^17^ Electron microscopy (EM) analysis for Httex1 showed a mixture of short fibrils and small oligomers in the early aggregation stages (time, 1 h), which explains the absence of a lag phase in the ThS assay. After longer incubation times (5 and 24 h), predominantly shorter fibrils with an average length of 190 ± 110 nm were observed as reported previously (Figure 5C).^17^ Interestingly, Htt171-43Q exhibited relatively slower aggregation kinetics with a lag phase of ∼ 2 h, and CD analysis indicated the presence of helical conformation in the monomeric Htt171-43Q, due to the helical rich region spanning residues 96-171 region, as discerned from the Cryo-EM of the full-length Htt protein.^31^ After 24 h, the Htt171-43Q CD spectrum showed a marked reduction in signal intensity (> 90%), suggesting both significant protein aggregation and precipitation (Figure 5B). The absence of a transition to β-sheet conformation in the CD spectra could be explained by the highly insoluble and compact nature of the aggregates formed by the Htt171-43Q protein compared to Httex1-43Q as evident from the EM studies of the same samples (Figure 5C).

**Figure 5.**
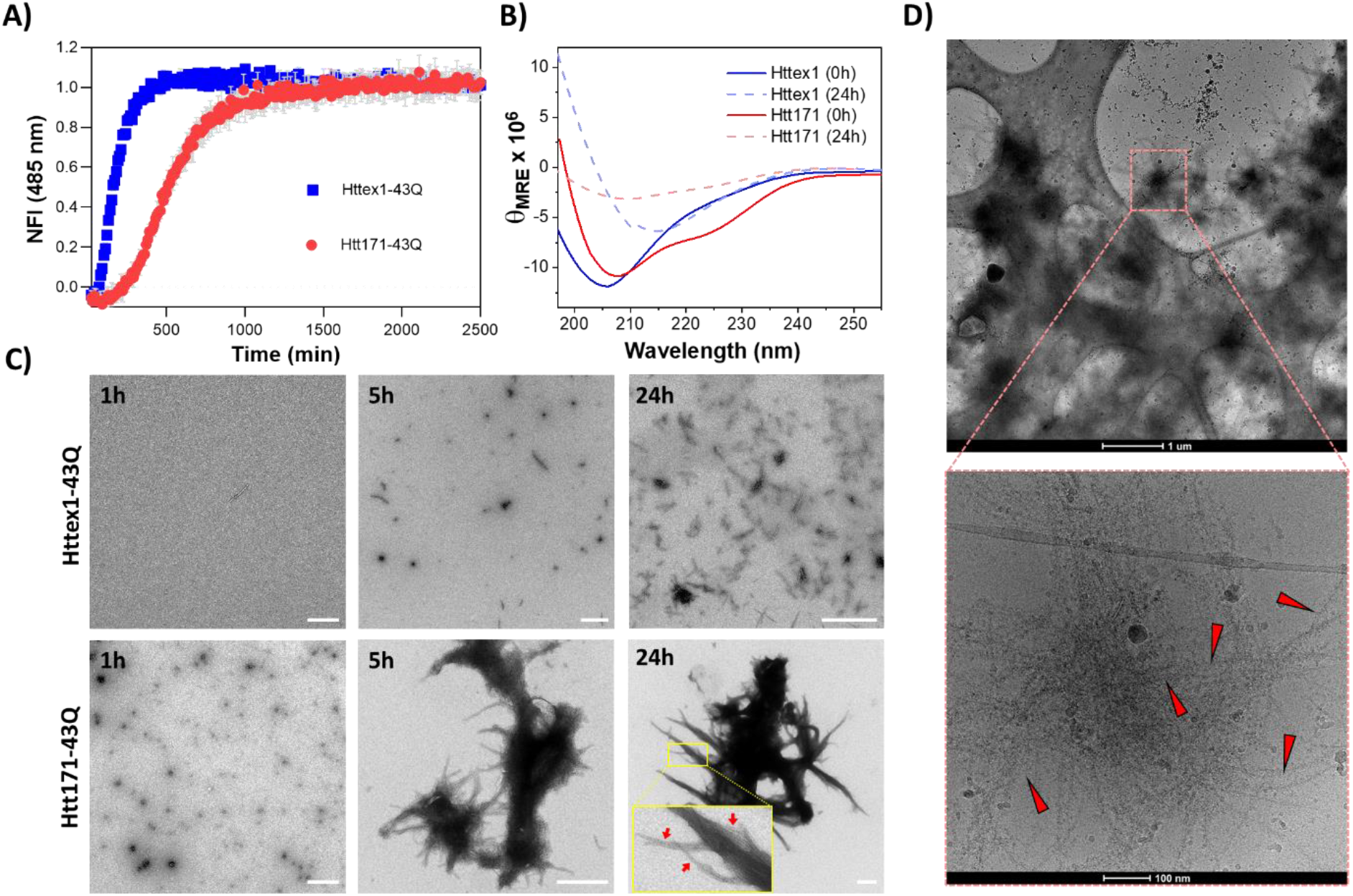
*In vitro* aggregation and structure of Httex1-43Q and Htt171-43Q proteins. A) Aggregation kinetics measured by ThS fluorescence at 485 nm (mean ± standard error of the mean [SEM], n=3). B) CD analysis of the proteins at 0 and 24h. C) Transmission electron microscopy (TEM) images of the proteins at 1, 5, and 24 h (scale bars are 500 nm). Red arrows indicate the fibrils inside the clumps. D) Cryo-EM images of Htt171-43Q. Red arrowheads represent fibrils.

In contrast to Httex1-43Q, we observed only oligomeric species at the early time point of Htt171-43Q aggregation, which explains the lag phase observed in the aggregation kinetics. However, at longer time points, these oligomers converted rapidly to form large clumps exhibiting fibril growth on their surface (Figure 5C). Due to the limitation of the negative staining EM analysis, it was not possible to examine the ultrastructural properties of Htt171-43Q in the dense regions of the clumps. Therefore, we performed Cryo-EM analysis, which clearly showed the presence of fibrils in the interior of the clumps, thus indicating the formation of fibrillar clumps by Htt171-43Q (Figure 5D). Despite several attempts to slow the aggregation reaction, we could never observe the formation of individual fibrils before the appearance of these large clumps of fibrillar aggregates. These observations clearly show differences in the aggregation mechanism observed for Httex1-43Q and Htt171-43Q and emphasize the role of structured domains beyond Httex1 (such as 96-17) in regulating the aggregation mechanism and morphology of the aggregates formed by N-terminal Htt fragments.

### Phosphorylation at T107 slows the aggregation kinetics of Htt171-43Q

To investigate the effect of site-specific phosphorylation at T107, S116, or both residues on the aggregation of wild type (23Q) and mutant (43Q) Htt171, we monitored the kinetics of aggregation of the unmodified and phosphorylated forms of these proteins (10 µM) using the ThS assay. The unmodified Htt171-23Q did not show any increase in ThS fluorescence over time, indicating that it did not aggregate or form amyloid-like fibrillar aggregates at this concentration (Figure S15). Similarly, phosphorylated Htt171-23Q (pT107, pS116, or pT107/pS116) proteins did not show any increase in ThS fluorescence. Transmission electron microscopy (TEM) images of these samples did not show any fibrillar aggregates for any of the Htt171-23Q proteins. However, a small percentage of amorphous aggregates were observed, and this observation explains the 20%– 25% loss of soluble protein observed for Htt171-23Q and phosphorylated Htt171-23Q during sedimentation assay (Figure S15). These findings demonstrate that WT and phosphorylated Htt171-23Q do not form fibrils *in vitro*.

In contrast, the unmodified Htt171-43Q showed an increase in the ThS fluorescence and reached the plateau in 14 to 16 h with a lag phase of 2 h (Figure 6A). Interestingly, phosphorylation of T107 caused a significant delay in the aggregation of Htt171-43Q with a longer lag phase (4.2h) and slower elongation phase (growth slope), whereas phosphorylation at S116 (Htt171-43Q-pS116) did not affect the lag phase of Htt171-43Q. Surprisingly, phosphorylation of T107 and S116 (Htt171-43Q-pT107-pS116) exhibited an increase in aggregation propensity of Htt171-43Q with a very short lag phase of 15 min. These results highlight the important role of cross-talk between the phosphorylated T107 and S116 residues with respect to modulating the aggregation of Htt171-43Q. Interestingly, when we monitored the aggregation using a sedimentation assay, where the extent of aggregation is assessed by monitoring the loss of soluble protein, we observed similar curves for Htt171-43Q, Htt171-43Q-pS116, and Htt171-43Q-pT107-pS116. However, Htt171-43Q-pT107 showed slightly slower kinetics, which was consistent with results from the ThS assay (Figure 6B).

**Figure 6.**
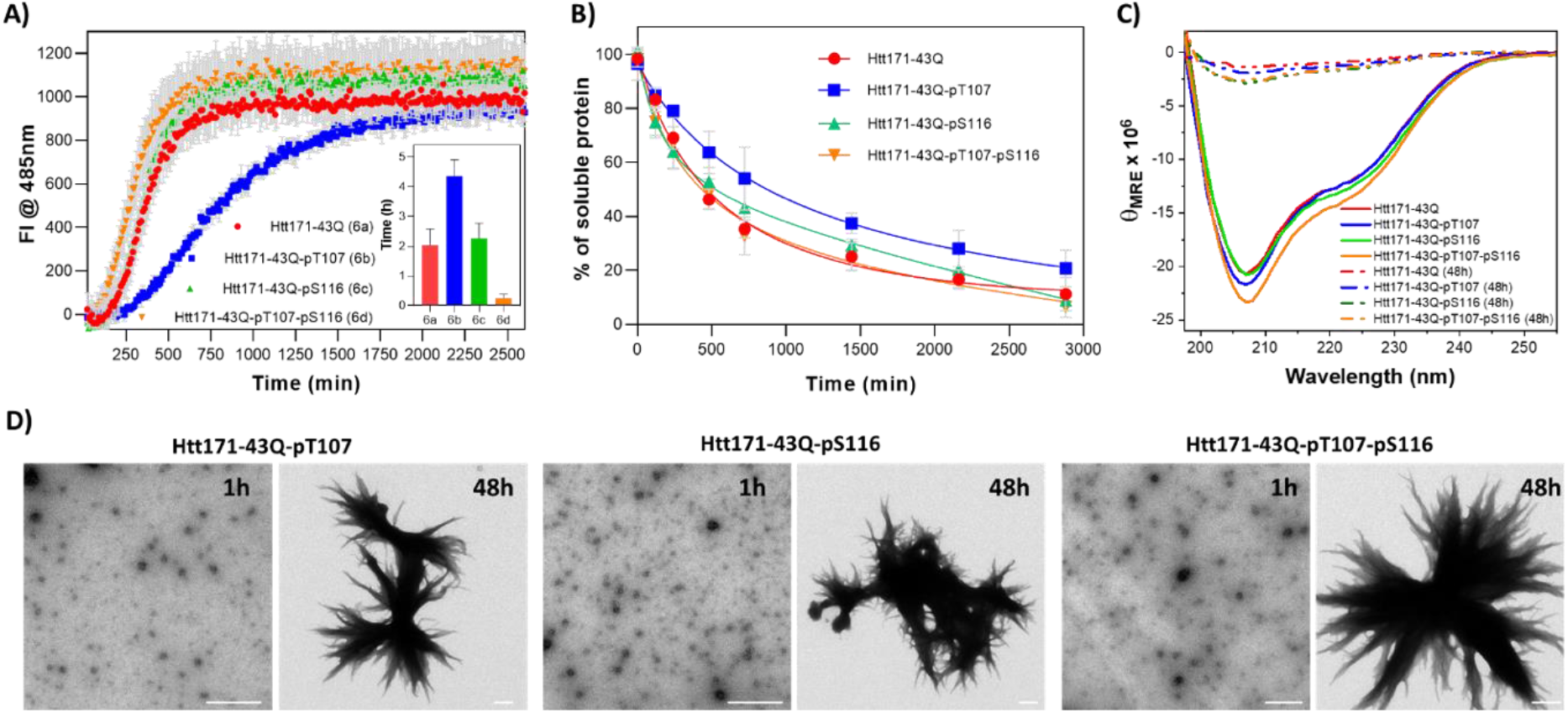
*In vitro* aggregation and structural properties of unmodified and phosphorylated Htt171-43Q proteins. (A) Aggregation kinetics measured by ThS fluorescence at 485 nm, insert: plot for lag phase of the Htt171-43Q proteins (mean ± SEM, n=3). (B) UPLC based sedimentation assay (loss of soluble protein) for the unmodified and phosphorylated Htt171-43Q (mean ± SEM, n=3). C) CD analysis of phosphorylated Htt171-43Q proteins at 0 and 48h. E) TEM images of the phosphorylated Htt171-43Q proteins at 1 and 48h (scale bars are 500 nm).

The CD spectra of the samples prior to initiating the aggregation reaction indicated that the unmodified and phosphorylated Htt171-23Q/43Q proteins adopted a predominantly helical conformation (Figure 6C), suggesting that phosphorylation at T107 or/and S116 does not significantly alter the conformation of Htt171-23Q/43Q. After 48 h, all samples showed > 90% reduction in intensity, suggesting loss of proteins due to precipitation, which is consistent with the high propensity of these proteins to form compact clumps of fibrillar aggregates (Figure 6D). Similar to Htt171-43Q, phosphorylated Htt171-43Q proteins show the presence of oligomeric species at an early time point (1 h), which transform to large fibrillar clumps at 48 h. No significant differences in oligomer or fibril morphology formed by Htt171-43Q and phosphorylated Htt171-43Q proteins were detected.

### Phosphorylation at T107 reduced the helicity of Htt105-171

Our results suggest that despite the presence of two highly aggregation-prone motifs in exons2 and 3, the aggregation of Htt171 is still driven primarily by the expanded polyQ repeat domain. However, phosphorylation of residues within 105-171 could still influence the kinetics and aggregation pathway of Httex1. Therefore, we sought to gain insight into the mechanisms by which this domain could modify the aggregation of Htt171. Given that this domain contains two aggregation-prone sequences, its propensity to aggregate and form fibrils *in vitro* was first assessed. We observed that neither unmodified nor any of the phosphorylated 105-171 fragments showed any tendency to aggregate or form amyloid-like fibrils as determined by the ThS assay and EM analysis (Figure S16). Furthermore, all four 105-171 fragments exhibited high stability to thermal denaturation and retained > 70% of their helical conformation upon heating to 90 °C and cooling back down to room temperature (Figure S16). These observations suggest that changes in the conformation of this domain or its interaction with other N-terminal domains, rather than its aggregation properties, could account for its role in modifying both the kinetics and the mechanism of Htt171 aggregation proteins. Therefore, we next sought to determine if phosphorylation at T07, S116, or both residues altered the conformation, thermodynamic stability, or aggregation properties of the structured domain corresponding to residues 105-171. We performed a CD analysis of unmodified and phosphorylated Htt105-171 fragments (**4a–c)**. The CD measurements revealed that the Htt105-171 protein adopted a predominantly α-helical conformation (Figure 7), consistent with the cryo-EM structure of Htt protein.^31^ The two phosphorylation sites, T107 and S116, are located in the helical domain and unstructured loop region of the Htt105-171 protein, respectively.

**Figure 7.**
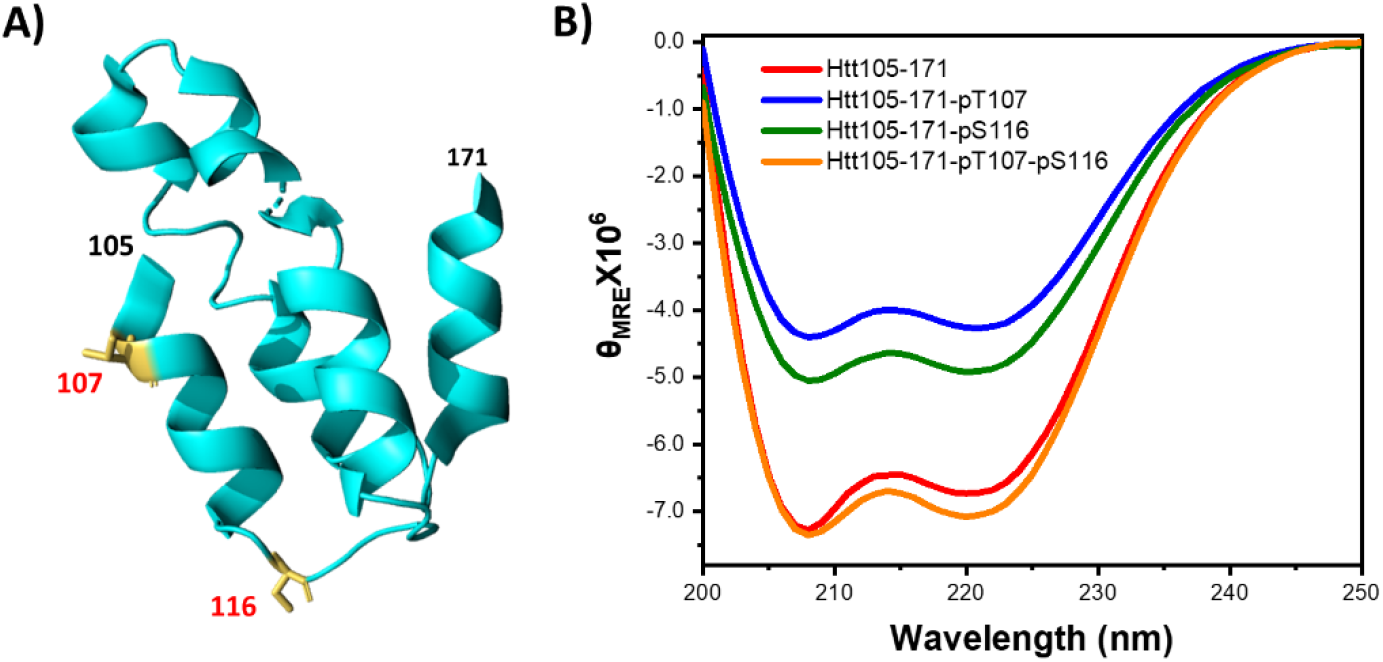
*In vitro* structure and aggregation of unmodified and phosphorylated Htt105-171 proteins. A) Rendered view of the secondary structure for Htt105-171 protein, the phosphorylation sites, T107 and S116 are highlighted yellow (PDB: 6EZ8). B) CD analysis of phosphorylated Htt105-171 proteins.

For Htt105-171, approximately 84% of its secondary structure existed in a α-helical conformation (Figure 7). However, introducing phosphorylation at T107 or S116 caused a reduction in the helicity to 54% and 64%, respectively, suggesting that phosphorylation at these residues destabilized the helical conformation of Htt105-171. Interestingly, Htt105-171 that was phosphorylated at both T107 and S116 exhibited a similar α-helical content (83%) as the unmodified peptide. The reduced helicity observed for Htt105-171-pT107 could probably explain the slower aggregation observed in the case of Htt171-43Q-pT107 and the reversal of this effect in the di-phosphorylated fragment (pT107/pS116) seemed to correlate with the reversal of the pT107-induced delay in aggregation in dephosphorylated Htt171 (pT107/pS116). Interestingly, this simple correlation was not applicable for phosphorylation at S116 in which reduction in the helicity of 105-171 did not translate into slower aggregation kinetics of Htt171-pS116. These findings suggest that helicity of this domain could play an important role in driving Htt171 oligomerization, but the effects of the helicity appear to be mediated by complex interactions with other domains in exon1. These interactions could be responsible for the formation of distinct types of oligomers with multiple fibril nucleation sites.

### The Htt171-43Q protein aggregates *via* distinct mechanism compared to Httex1

Our results on unmodified and phosphorylated Htt171 reveal that this longer fragment formed fibrils with distinct morphological properties, suggesting a different aggregation mechanism compared to mutant Httex1. Therefore, we sought to monitor the evolution of the aggregation process by performing time-dependent EM studies. Immediately after the initiation of Htt171-43Q aggregation, we observed the formation of large oligomers (Figure 8) with a mean diameter of 46 ± 25 nm (n = 60), which were much larger than what we and others previously observed for prefibrillar oligomers of Httex1-43Q.^34^ At later time points (10–15 min), we observed that amorphous oligomers tended to agglomerate and formed clusters of oligomers that seemed to align in ordered structures. A closer look at these structures showed the presence of what appeared to be nucleation sites for fibril formation and growth on their surfaces (Figure 8A). Interestingly, several nucleation sites were observed, suggesting an independent nucleation site for fibrillar growth. Although mutant Httex1 has also been shown to form globular oligomeric species during the initial stage of aggregation,^35^ the size of these oligomers is much smaller (mean width of 87 ± 25 nm) than the size we observed for Htt171-43Q (ranging between 280 and 820 nm). At 60 and 90 min, the presence of fibrils growing from the surfaces of oligomers became more evident. The fibrils were long, ranging from 500 nm to 1 μM in length and exhibited high lateral association in contrast to short and dispersed fibrils as observed for Httex1-43Q (Figures 8A and S17). These observations indicate a significant difference in the mechanism of aggregation and morphology of the fibrils formed by Htt171-43Q and Httex1-43Q. At longer time points (> 6 h), only the presence of large fibrillar clumps with very few or no globular oligomers was noted. To further validate our EM observations, a time-dependent atomic force microscopy (AFM) analysis of Htt171-43Q (Figure 8B) was performed. Similar to our EM data, oligomer formation at early stages (10 min) of aggregation were observed, and at longer time points (60 min), fibrils growing from the surfaces of oligomers were then observed. Based on the time-dependent EM and AFM analysis, we hypothesized an aggregation mechanism in which monomers aggregate to form oligomers of various size, possibly driven by phase separation. These oligomers agglomerate to form large globular species that serve as nucleation sites for initiating fibril growth. Given that these fibrils grow from multiple nucleation sites on the oligomers, they exhibited high tendency to self-associate resulting in the formation of tightly packed clumps of fibril assemblies (Figure 8E). Our observations suggest that Htt171-43Q possesses a more complex aggregation mechanism in which the structured C-terminal domain (Htt96-171) plays an important role in regulating early events in the aggregation process of Htt171-43Q.

**Figure 8.**
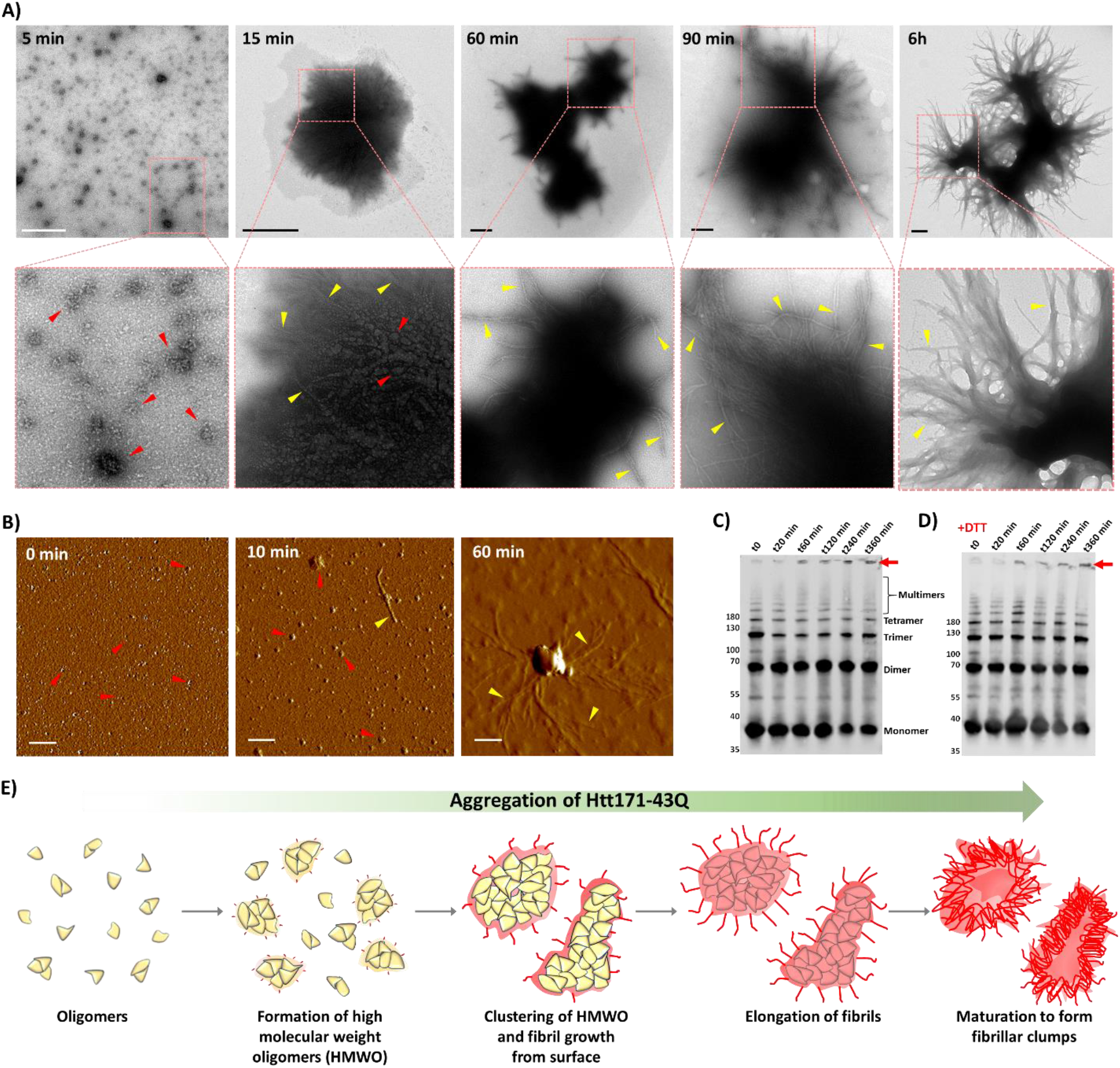
Studying the aggregation mechanism of Htt171-43Q. Time-dependent (A) TEM analysis and (B) AFM analysis for monitoring the aggregating Htt171-43Q protein. C) Western blot for the aggregating Htt2-171-43Q at various time points in the (C) absence and (D) presence of reducing agent (DTT), red arrow indicates the high molecular aggregates accumulated in the stacking gel. E) Schematic representation of the different events occurring during aggregation based on the observations made from TEM, atomic force microscopy (AFM), and Western blot analysis (scale bars are 500 nm for the EM and AFM micrographs).

To further investigate early oligomerization events and assess the aggregation state of Htt171 oligomers that precede the formation of fibrils, we assessed their size distribution by Western blotting. Figure 8C and D show the presence of the formation of dimer, trimer, tetramer and higher-order multimers (Figure 8C) as early as the first 10 min after sample incubation. Over time, we observed an increase in the formation of higher molecular weight aggregates that accumulated in the stacking gel. Similar results were observed in samples containing reducing buffers, indicating that the high molecular weight species formation is not driven by cysteine (C105, C109, C137 and C152)-based intermolecular disulfide bond formation (Figure 8D). These observations indicate that Htt171-43Q underwent fast oligomerization even in the presence of a reducing agent (dithiothreitol).

Finally, to determine which region in the C-terminal domain could be responsible for driving the aggregation of Htt171-43Q *via* the pathway leading to the formation of these tightly packed fibrillar clumps, we performed aggregation studies and EM analysis of shorter N-terminal Htt fragments, Htt104-43Q and Htt140-43Q. Interestingly, the Htt104-43Q fragment formed shorter and well-separated single fibrils, similar to those observed for Httex1-43Q, whereas the Htt140-43Q formed large clumps with laterally associated fibrils growing on their surfaces, similar to what was observed for Htt171-43Q (Figure 9). Htt140-43Q exhibited a shorter lag phase and faster aggregation kinetics in comparison to Htt171-43Q (Figure 9). These observations point to the C-terminal structured domain (Htt105-140), which encompasses the two highly aggregation domains (Htt105-116 and Htt128-138) that were previously identified by our group, as being responsible for the aggregation properties observed for the longer N-terminal fragments Htt140 and Htt171.^15^

**Figure 9.**
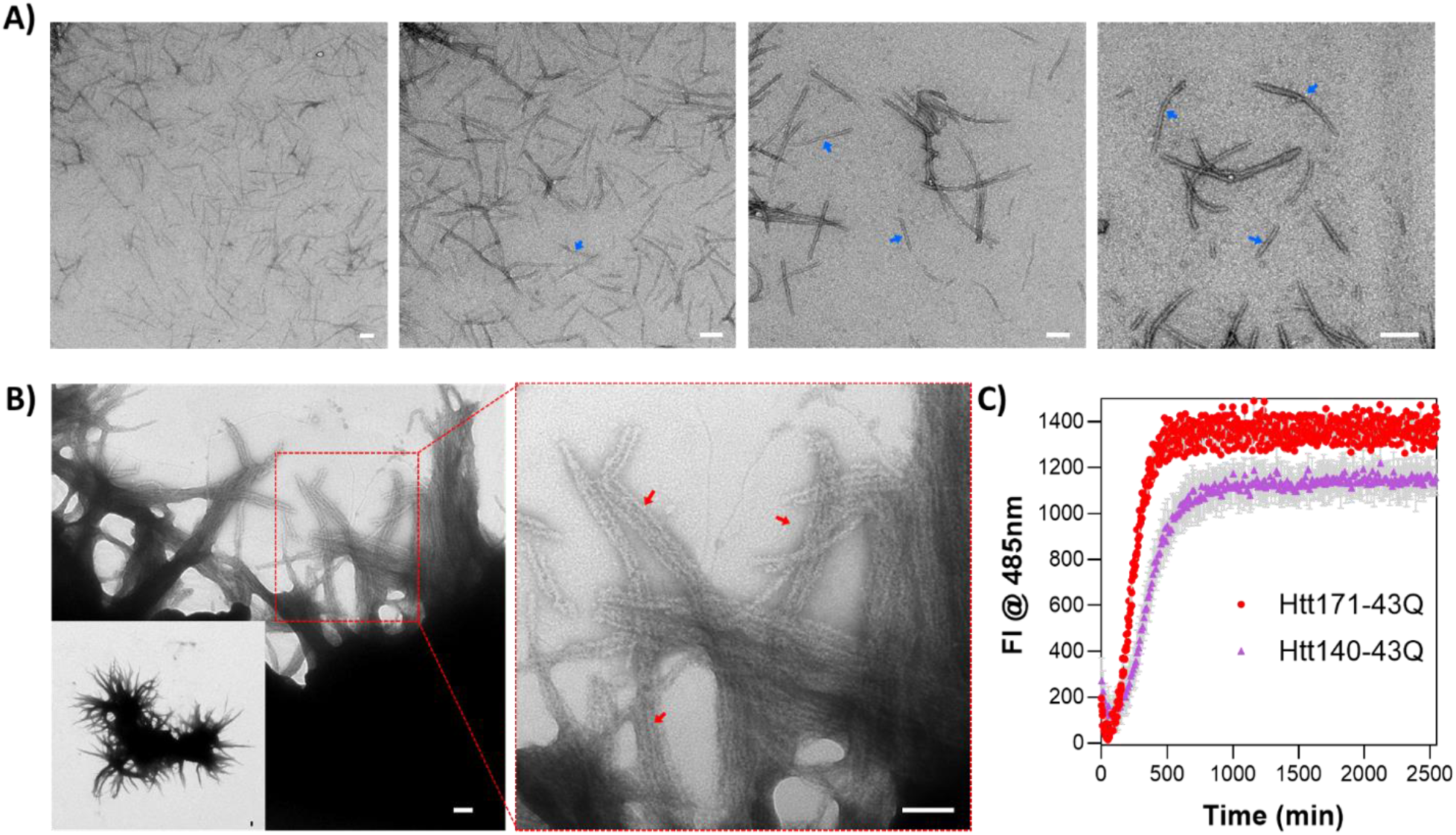
A) TEM images of Htt104-43Q-Mesna (10 µM) protein after 24h of aggregation. Blue arrows indicate the fibrils. B) TEM images of Htt140-43Q (10 µM) after 24h of aggregation. Red arrows indicate the fibril growth on surface and lateral association (scale bars are 100 nm for all the micrographs). C) Aggregation kinetics measured by ThS fluorescence at 485 nm (mean ± SEM, n=3).

## Discussion

Several studies have shown that overexpression of mutant Httex1 with expanded polyQ repeats (43-150Q) recapitulates many features of HD pathology formation and symptomology in several cellular and *in vivo* models.^3^ These observations made Httex1 among the most studied N-terminal fragment in HD models and the main Htt fragment used in *in vitro* aggregation studies to investigate mechanisms of Htt aggregation.^5^ However, it has been well-established that other longer N-terminal polyQ-containing fragments can be detected under both physiological and pathogenic conditions.^36,28^ Mutant forms of several longer N-terminal fragments, such as Htt118, Htt171, Htt536, and Htt586, have also been found in Htt inclusions.^37,26,38,30^ Among them, the Htt171 fragment is one of the most studied fragments in cellular and *in vivo* models of HD.^28,39,26^ Despite this finding, no reports concerning *in vitro* structural and aggregation properties of Htt171 or any N-terminal Htt fragments longer than Httex1 exist in the literature. Herein, we present a robust and efficient protein expression and purification method for producing mg quantities of wild type (WT/23Q) and mutant (43Q) N-terminal fragments, including Htt171, Htt140, and Htt1-104 and new chemoenzymatic semi-synthetic strategies that for the first time allow site-specific phosphorylation beyond exon1. These advances have been facilitated by the discovery of novel kinases that phosphorylate Htt at T107, S116, and both residues, thus enabling us to gain novel insight into how sequences and PTMs beyond exon1 influence the aggregation mutant Htt fragments of various lengths.

Recent studies have identified several novel phosphorylation sites within exon2 and exon3 of the Htt protein and suggest that blocking phosphorylation at S116 may attenuate Htt toxicity.^17^ However, these studies were primarily based on the mutations that block or mimic phosphorylation and did not investigate the effects of these mutations on Htt aggregation or inclusion formation. Therefore, the mechanisms by which phosphorylation at these residues influence mutant Htt aggregation and toxicity of longer N-terminal Htt fragments remains unknown. Interestingly, several of the newly identified phosphorylation sites occur in close proximity to previously identified putative cleavage sites that can lead to the generation of shorter N-terminal fragments.^13,14^ To address this knowledge gap, we developed new chemoenzymatic methods to generate WT and mutant Htt171 proteins that are site-specifically phosphorylated at single (pT107 and pS116) or multiple (pT107/pS116) sites and then investigated their conformation and *in vitro* aggregation properties. Our findings show that phosphorylation at S116 did not affect the aggregation and the morphology of the aggregates formed by Htt171, whereas phosphorylation at T107 significantly slowed its aggregation. Interestingly, phosphorylation at T107 and S116 accelerated the aggregation of Htt171. This observation in combination with previous observations from our group on the cross-talk between T3 phosphorylation and K6 acetylation underscores the importance of conducting further studies to elucidate the role of cross-talk among PTMs in the regulation of Htt aggregation, especially since several PTMs are known to cluster in a short sequence motifs throughout the Htt protein (Figure 1a).^17,40^ For example, most of the proteolytic sites identified on Htt are flanked by phosphorylation residues, indicating possible cross-talk between phosphorylation and proteolysis.^6^

Although several studies have investigated the aggregation propensity of various N-terminal Htt fragments in different cellular and animal HD models, characterization of the aggregation properties of these fragments was limited to quantifying the number of inclusions and their size or shape.^41^ These studies demonstrated N-terminal length-dependent changes in these properties in cells but did not offer insight into the molecular and structural basis underlying these differences. In this study, we investigated and compared the structure and aggregation properties of longer N-terminal Htt fragments (Htt171-43Q, Htt140-43Q, and Htt104-43Q). Our findings demonstrate that mutant Htt171 aggregate via a distinct aggregation mechanism compared to Httex1. Their aggregation properties are characterized by 1) a more extended lag phase; 2) transient population of large disordered oligomers, and 3) the formation of tightly packed fibrils that emanate from a central core (Figure 8).

These findings suggest that the presence of the C-terminal structured domains beyond Httex1 (e.g., 96-171) significantly alters the aggregation kinetics and pathway of Htt N-terminal fragments. On the basis of these findings, we proposed a model of aggregation for Htt171 that is initiated by C-terminal structured domain through a strong intermolecular coil–coil interaction, which leads to the formation of a-helix rich oligomeric species (Figure 10A). This oligomerization causes a rise in the local concentration of polyQ and enhances intra- and inter-molecular polyQ associations. Conformational reorganization will occur within the oligomers, and multiple nucleation sites β-sheet rich would be formed. The nucleation sites will then recruit Htt171 monomers, a process leading to the formation of fibrils that emanate from multiple site on the surface of these oligomers. Interestingly, ataxin-3, a polyglutamine protein associated with neurodegenerative disorders has been reported to exhibit a similar multistep aggregation mechanism^42^ in which the N-terminal structured domain of the protein initiate association of monomers to form oligomers followed by polyQ-dependent aggregation. This model differs from those that have been proposed for the mechanism of mutant Httex1 oligomerization and fibril formation. Previously, it was suggested that the helical propensity of the Nt17 domain plays a key role in initiating the fibrillization of mutant Httex1 by driving early oligomerization events that bring the polyQ domains in close proximity with each other, thus increasing their effective concentration. The transition from oligomers to fibrils is driven by the transition of the polyQ domain to β-sheet conformation, which then dominates as the main driver of the fibrillization process (Figure 10B).^43,44,45,46^ An alternative model has been proposed in which early oligomerization events are driven by the formation of a more globular amyloidogenic intermediate when the Nt17 interacts with the polyQ domain (Figure 10C).^47,48^ Further studies are underway to evaluate our hypothesis and further elucidate the sequence, structural determinants, and mechanisms of longer N-terminal Htt proteins.

**Figure 10.**
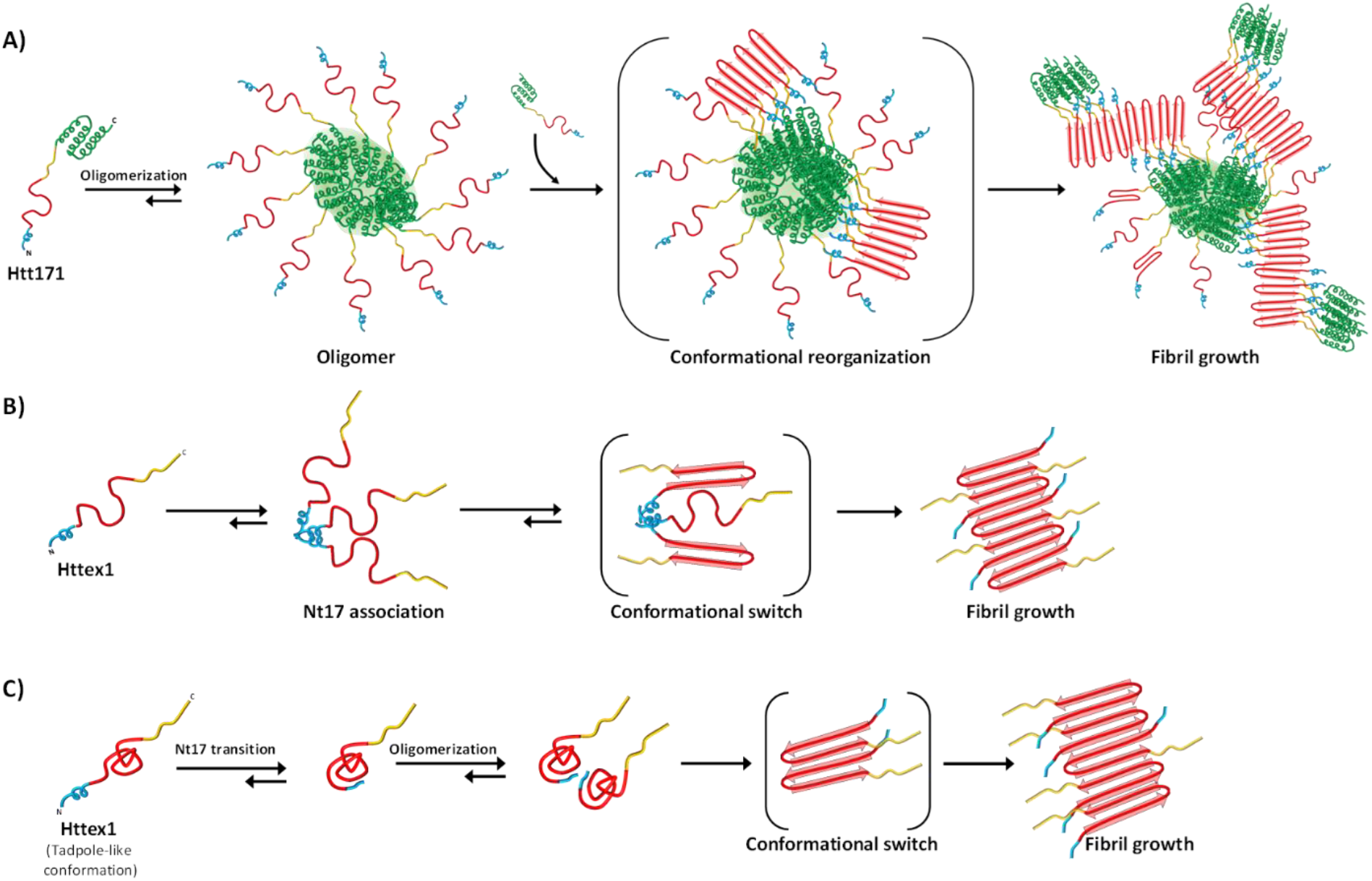
A schematic model for aggregation of Htt171 and Httex1. A) We propose that the initial step for the aggregation of Htt171 (Nt17, blue; polyQ, red; proline rich domain, yellow; green, C-terminal region) is the association of the C-terminal structured domain to form oligomers. These oligomers undergo conformational reorganization and form polyQ driven β-sheet rich nucleation sites. Then the elongation of the nuclei by recruiting monomers generates fibrils. B) The disordered monomeric Httex1 spontaneously assemble into an oligomer formation which is mediated by the association of the α-helical conformation of Nt17. Oligomerization results in a high local concentration of polyQ and subsequent conformational switch of polyQ in β-sheet structure. Elongation of nuclei by monomer addition generates amyloid fibrils. C) The Httex1 acquires a tadpole like conformation in the monomeric form. The Nt17 interacts with the polyQ, which enhances intermolecular hydrophobic interactions and facilitates oligomerization. High local concentration of polyQ in oligomers cause conformational switch to β-sheet structure, followed by fibril growth.

Consistent with these observations, a previous study by Moore *et al*. demonstrated distinct morphology of the inclusions for the N-terminal Htt fragments of less than 171 (Httex1, Htt105, and Htt117) compared to longer Htt fragments (Htt171, Htt513, Htt536, and Htt586) in a cellular model of Htt inclusion formation.^41^ Htt105 and Htt117 formed larger inclusion and exhibited poly-Q dependent aggregation properties, whereas Htt171 formed small puncta that were scattered throughout the cytoplasm. The longer N-terminal fragments (Htt536 and Htt586) formed a mixture of small and larger inclusions irrespective of the polyQ-repeat length. In a more recent report, Anjalika *et al*. showed that while both mutant Httex1 and Htt171form cytoplasmic inclusions, only mutant Httex1 was capable of inducing toxicity in a Drosophila model of HD^37^, suggesting that the length of the N-terminal fragments dramatically influences the toxicity of the protein. Interestingly, in this model, the addition of 28 AAs to Httex1 (Htt118) caused a reversal of the aggregation and toxicity of the expanded polyQ repeat, whereas the addition of 81 C-terminal AAs (Htt171) restored the aggregation. These observations along with our *in vitro* studies demonstrate that different N-terminal fragments could have distinct mechanisms of aggregation and toxicity. We hypothesize that the sequences outside the polyQ domain or cross-talk between the polyQ domain and structured domains in another part of Htt could play important roles not only in the initiation of Htt aggregation but also in determining the Htt aggregation pathway and final morphological or structural properties of the aggregates. Given that several polyQ containing N-terminal fragments have been reported to coexist in the HD brain and Htt inclusions, it is crucial to revisit the human pathology to define the exact sequences of these fragments and their relative contribution to inclusion formation. It is plausible that the nature of the dominant aggregating fragment in HD inclusions varies among individuals with the same or different polyQ expansions. Such variations could partially explain the large variations in age of onset among HD patients with the same polyQ repeat lengths. Studies are underway in our laboratory to investigate the cross-talk among different Htt N-terminal fragments and how those fragments influence another one’s aggregation.

### Therapeutic implications and Conclusion

Previous strategies aimed at inhibiting mutant Htt aggregation have focused primarily on targeting polyQ dependent aggregation events or PTMs within the Nt17 domain. Therefore, most Htt *in vitro* aggregation studies and drug screening efforts to identify inhibitors of Htt aggregation are mostly based on poly-Q peptides, mutant Httex1, or mutant Httex1 fused to other peptides/proteins as substrates. Although these efforts have led to the identification of several small molecules (such as Niclosamide, AMG 9810, Methylene blue, PGL-135, and arginine ethyl ester) that inhibit the aggregation of polyQ peptides and Httex1, no aggregation inhibitors have advanced to clinical trials.^49,50,51,52^ The observed differences in the aggregation mechanism of the longer N-terminal Htt fragments further emphasize the prominent role of cross-talk between the polyQ repeat domain and other flanking C-terminal sequences in determining the Htt aggregation pathway and the ultrastructural properties of the Htt aggregates. Therefore, we propose that future screening efforts to identify Htt aggregation inhibitors should be based on multiple physiologically and pathologically relevant polyQ-containing fragments of different lengths. This approach is more likely to lead to the identification of anti-aggregation drugs capable of blocking the aggregation of multiple Htt N-terminal fragments and not just Httex1, which is only one of the fragments found in Htt inclusions.

The phosphorylation sites, T107, S116, and S120, that were investigated in this study form a phosphorylation cluster in the region 81-129 in which several putative cleavage sites have been reported.^14^ Cleavage within this region would result in the generation of shorter N-terminal fragments that are more aggregation-prone and toxic (such as Httex1). Phosphorylation at S434 by cyclin-dependent kinase (Cdk5) has been shown to reduce cleavage by caspase-3 of Htt.^53^ Similarly, mimicking phosphorylating at S536 inhibited calpain cleavage of S536 and subsequently led to a reduction in Htt toxicity.^10^ Therefore, we hypothesize that the T107 and S116 phosphorylation sites could play important roles in regulating the proteolytic cleavage and generation of toxic N-terminal Htt fragment. Our ability to generate pure unmodified and phosphorylated N-terminal fragments, such as Htt140 and Htt171, should facilitate future studies to identify the enzymes responsible for cleaving Htt and to investigate the cross-talk between phosphorylation and proteolysis in this region. If increasing or reducing phosphorylation at these residues enhances or blocks Htt proteolysis and generation of the shorter and more toxic fragments, this finding would suggest that targeting the kinases that regulate phosphorylation at these residues could offer an alternative strategy to prevent Htt aggregation and toxicity.

## Supporting information

Raja et al. supporting information

## Associated content

### Supporting information

Materials and methods, expression, purification and characterization of proteins (Htt105-171, Htt2-104-23Q/43Q-Mesna and Htt2-171 23Q/43Q), SPPS, kinase screening, site-specific phosphorylation of Htt105-171, semi-synthesis of unmodified and phosphorylated Htt171-23Q/43Q, aggregation kinetics, CD analysis, EM and AFM analysis and supporting data.

## Acknowledgment

This work was supported by CHDI and École Polytechnique Fédérale de Lausanne. We thank Dr. Senthil Kumar Thangaraj, and Dr. Pedro Santana Magalhães, EPFL, for critical review of the manuscript.

