## Supplementary material for "A new chemoenzymatic semisynthetic approach provides novel insight into the role of phosphorylation beyond exon1 of Huntingtin and reveals N-terminal fragment length-dependent distinct mechanisms of aggregation": Raja et al. supporting information

### Table of content

1. Materials and Methods……………………………………………………………….3

2. Expression and purification of Htt105-171 fragment……………………………………...4

3. Expression and purification of Htt2-104-23Q/43Q-Mesna…………………………………..4

4. Expression and purification of Htt2-171 23Q/43Q…………………………………………..5

5. Expression and purification of Htt105-171-T107A, Htt105-171-S116A and………………..6
Htt105-171-S120A.

6. Design strategy and SPPS of phosphorylated Htt105-171…………………………………...6

7. Dot blot analysis to validate Anti-pT107 and Anti-pS116 antibodies………………………..8

8. Kinase screening for identifying site-specific kinases……………………………………….8

9. Determining the NLK phosphorylation site on Htt105-171………………………………….9

10. Semi-synthesis of Htt105-171-pT107, Htt105-171-pS116, and Htt105-171-pT107-pS116..9

11. Semi-synthesis of phosphorylated Htt171-23Q/43Q1…………………………………….10

18. Reference……………………………………………………………………………………….24

### 1. Materials and Methods

All of the proteins used in these studies were produced, purified, and characterized in-house. Trifluoroacetic acid (TFA), formic acid (LC-MS grade), sodium 2-mercaptoethanesulfonate (Mesna), Tris(2-carboxyethyl)-phosphine hydrochloride (TCEP), exafluoroisopropanol (HFIP), magnesium chloride ($MgCl_2$), thioflavin T (ThT) 4- mercaptophenylacetic acid (MPAA), L-proline, and D-trehalose were purchased from Sigma-Aldrich. HPLC-grade acetonitrile was purchased from Merck or Applied Biosciences. Dimethyl sulfoxide (DMSO), dithiothreitol (DTT), isopropyl β–D-1- thiogalactopyranoside (IPTG), phenylmethanesulfonyl fluoride (PMSF), and ethylene glycol-bis(β-aminoethyl ether)-N,N,N′,N′-tetraacetic acid (EGTA) were purchased from Applichem. Adenosine triphosphate (ATP) was purchased from Genbiotech. Secondary goat or rabbit anti-mouse antibody labelled with Alexa Fluor 680 were purchased from Invitrogen. The fusion proteins were purified on fast performance liquid chromatography (FPLC) from GE. Preparatory and semi-preparatory high-performance liquid chromatography (HPLC) was performed on a Waters 2535 HPLC system using a C4 column (Phenomenex Jupiter C4 or Higgins, 10 μm, 300Å, 21.2 x 250 mm and 5 μm, 300Å, 10 x 250 mm) with ultraviolet (UV) detection at 214 nm and manual fraction collection. Liquid chromatography/mass spectrometry (LC/MS) was performed with a Thermo LTQ instrument, and the ESI technique was used for ionization. Samples were run through a C3 column (1 x 75 mm, 5 μm) with a gradient of 5% to 95% acetonitrile at 0.3 mL/min over a period of 10 min. UPLC analysis was performed on a Waters Acquity H-Class system using a C4 column (1.7 μm, 300Å, 2.1 x 150 mm) with UV detection at 214 nm and run time of 3 min (gradient 10% to 90% acetonitrile) with 0.6 mL/min flow rate. For negative stain TEM, Formvar carbon film on 200-mesh grids and uranyl formate from Electron Microscopy Sciences were used. Circular dichroism (CD) measurements were performed in a Chirascan Circular Dichroism Spectrometer by Heidi Esbensen from Applied Photophysics. A prestained protein ladder (PageRular, 26617) was purchased from Thermo Scientific. The nitrocellulose membrane was purchased from Thermo Fisher Scientific for dot and western blot analyses. Dot and western blots were scanned using the Odyssey® CLx System. The antibodies to detect phosphorylation at T107 and S116 were generated by Eurogentec using synthesised Htt101-113-pT107 and Htt113-125-pS116 as antigens to induce immunization and generate antibodies.

**2. Expression and purification of Htt105-171 fragment**

The expression of the construct was performed in chemo-competent *E. coli* ER2566 cells. The bacterial transformation was performed with pTWIN1-Htt105-171-Sumo-$His_6$ plasmid and the transfected cells were grown in Luria Bertani (LB) media (with 100mg/mL ampicillin) at 37 ºC until an $OD_{600}$ of 2.5 was reached. The preculture media was diluted to 0.05 at $OD_{600}$ and grown at 37 °C (180 rpm) until it reached an $OD_{600}$ of 0.5. The culture media was transferred to an 18 ºC incubator. Expression was then induced with 1 mM IPTGand incubated for 18 h. Bacteria were harvested by centrifugation, resuspended in lysis buffer (50 mM Tris, 500 mM NaCl, 15 mM imidazole, pH adjusted to 7.5, 0.3 mM PMSF, and protease inhibitor tablets), and lysed by ultrasonication. The supernatant was separated from the cell debris by centrifugation (23,000 x g, 50 min, 4 °C). The Htt105-171-SUMO-$His_6$ fusion protein was purified from the bacterial lysate by nickel-immobilized metal affinity chromatography (Ni-IMAC) on an FPLC system using several steps (Figure S2). The bacterial lysate was loaded onto the Ni-NTA column, washed with buffer A (50 mM Tris, 500 mM NaCl, 15 mM imidazole, pH adjusted to 7.5), and the fusion protein was eluted with buffer B (50 mM Tris, 500 mM NaCl, 500 mM imidazole, pH adjusted to 7.5). The eluted fractions were analyzed using SDS PAGE to determine the fractions containing the fusion protein after which these fractions were pooled accordingly (Figure S2). DTT powder was directly added and dissolved to a final concentration of 100 mM. Finally, ubiquitin-like-specific protease 1 (ULP1) was added and incubated at 4 ºC for 14 h to cleave the SUMO-$His_6$. The Htt105-171 was purified by preparative HPLC with a gradient from 35% to 75% acetonitrile in $H_2O$ with 0.1% TFA over 50 min. The desired protein eluted at approximately 52 min, and the collected HPLC fractions were analyzed using ESI-MS and UPLC. Pure fractions were pooled and lyophilized to yield a white fluffy lyophilizate (approximately 6.0 mg/L).

**3. Expression and purification of Htt2-104-23Q/43Q-Mesna**

Bacterial transformation was performed with pTWIN1-Htt2-104-23Q/43Q-Intein-$His_6$ plasmid in *E. coli* ER2566 cells, and the transfected cells were cultured, lysed, and purified though Ni-IMAC on FPLC (as mentioned in the expression and purification of Htt105-171). The eluted fractions from FPLC were analyzed using SDS-PAGE to identify the fractions containing the fusion protein and pooled accordingly (Figure S1). Intein splicing and thioester formation were induced by adding Mesna to a final concentration of 100 mM in the pooled fractions and incubated at room temperature for 14 h (23Q) or 37 ºC for 2 h (43Q). Cleavage was monitored through ESI-MS to determine the completion of the reaction. The Htt2-104-Mesna proteins were purified via preparative HPLC with a gradient ranging from 25% to 40% acetonitrile in water over 50 minutes. The desired protein eluted approximately at 35 min (23Q) or 33 min (43Q), and the collected HPLC fractions were analyzed with ESI-MS and UPLC (Figure S1). Pure fractions were pooled and lyophilized to yield a white fluffy lyophilizate (approximately 3.36 and 0.96 mg/L for Htt2-104-23Q-Mesna and Htt2-104-43Q-Mesna, respectively). The higher aggregation propensity of Htt2-104-43Q-Mesna caused a significant decrease in the yield. The expression and purification of Htt2-104-43Q-Mesna presented a series of challenges, which required extensive optimization of several parameters of expression: (1) shortening the expression time to 14 instead of 18 h; (2) lowering the temperature to 16 °C; 3) performing the Ni-IMAC purification, intein cleavage, and HPLC purification continually; and 4) keeping the protein on ice at all time to prevent aggregation.

**4. Expression and purification of Htt2-171 23Q/43Q**

Bacterial transformation was performed with pTWIN1-Htt2-171-23Q/43Q-Sumo-$His_6$ plasmid in *E. coli* ER2566 cells, and the transfected cells were cultured, lysed, and purified with Ni-IMAC on an FPLC machine (as mentioned in the expression and purification of Htt105-171). The eluted fractions from FPLC were analyzed using SDS-PAGE to identify the fractions containing the fusion protein and were then pooled accordingly. DTT powder was directly added and dissolved to a final concentration of 100 mM (Figure S3). Finally, ULP1 was added and incubated at 4 ºC for 14 h (23Q) or 30 min (43Q, cleavage was monitored through UPLC to determine the completion of the reaction) to cleave SUMO-$His_6$. The Htt2-171 was purified by preparative HPLC with a gradient ranging from 15% to 70% acetonitrile in water over 50 min for Htt2-171-23Q and 40% to 70% acetonitrile in water over 50 min for Htt2-171-43Q. The desired protein eluted at approximately 46 min (23Q) or 50 min (43Q), and the collected HPLC fractions were analyzed via ESI-MS and UPLC (Figure S3). Pure fractions were pooled and lyophilized to yield a white fluffy lyophilizate (approximately 4.88 and 1.99 mg/L for Htt2-171-23Q and Htt2-171-43Q, respectively).

**5. Expression and purification of Htt105-171-T107A, Htt105-171-S116A and Htt105-171-S120A**.

Good quantities of the recombinant proteins were expressed and purified following the Htt105-171 expression and purification protocol (approximately 3.80, 5.4, and 4 mg/L for Htt105-171-T107A, Htt105-171-S116A, and Htt105-171-S120A, respectively, Figure S4).

### 6. Design strategy and SPPS of phosphorylated Htt105-171

We designed a two-fragment strategy to access site-specific phosphorylation of Htt105-171 fragments. To maintain the native sequence, a native cysteine residue (Cys138) was selected as the ligation site. The two fragments, Htt105-137 (with site-specific phosphorylation at T107, S116, or S120) were synthesized through SPPS, and Htt138-171 would be generated using a bacterial expression system (Figure S6). Initially, we attempted to synthesize Thz-Htt106-137-Nbz-pS116 via Fmoc-solid phase peptide synthesis (Fmoc-SPPS) using a Fmoc-MeDbz (3-Fmoc-amino-4-(methylamino)benzoic acid)-functionalized Rink amide resin (0.25 mmol/g) and 1-[Bis(dimethylamino)methylene]-1H-1,2,3-triazolo[4,5-b]pyridinium 3-oxide hexafluorophosphate, Hexafluorophosphate Azabenzotriazole Tetramethyl Uronium (HATU) as a coupling reagent. During our initial efforts, we faced difficulties in proceeding with synthesis beyond the Leu126 residue, which may have been due to the presence of multiple hydrophobic amino acids (AAs) as shown in Figure S7. To overcome this challenge, we changed the solvent from dimethylformamide (DMF) to N-methyl-2-pyrrolidone (NMP) and performed double coupling for all of the AAs. Under these modified conditions, the quality of the synthesis showed a significant improvement. Furthermore, pseudo-proline dipeptide was introduced at Asn120-Ser121 to increase peptide solvation and improve the quality of the synthesis. Phosphorylation at the S116 residue was introduced using Fmoc-Ser(PO-(OBzl)OH)-OH. Beyond the S116 AAs, coupling efficiency was drastically reduced; to overcome this challenge, we performed further AA couplings in the microwave peptide synthesizer. The synthesis of the Thz-Htt106-137-Nbz-pS116 was complete; however, the synthesis quality was poor. Moreover, the crude product had very low solubility, which hindered the purification of the peptide. The insoluble nature of the synthesized Htt105-138 fragment could be explained by the presence of two hydrophobic aggregation domains (Htt108-116 and Htt128-138). Similarly, we dedicated our efforts to synthesizing Thz-Htt106-137-Nbz-pT07; however, poor synthetic quality and insolubility hindered our access to the phosphorylated fragment (Figure S8).

**6.1 Synthesis of Thz-Htt106-137-Nbz-pS116**. Thz-Htt106-137-Nbz-pS116 was synthesised on Rink amide resin (0.25 mmol/g, 0.1 mmol scale) according to the following procedure (Figure S7). Initially resin was coupled (double coupling) with unnatural amino acid, Fmoc-Dbz-OH manually for 1 h using 4 eq of 1- [bis(dimethylamino)methylene]-1H-1,2,3-triazolo[4,5-b] pyridinium 3-oxid hexafluorophosphate (HATU) and 6 equiv of N,Ndiisopropylethylamine (DIEA). The resin was then transferred to a peptide synthesizer (336X peptide synthesizer from CS Bio) for further synthesis. Double coupling for all of the AAs with a coupling time of 45 min each was performed in DMF or N-methylpyrrolidone (NMP). Deprotection was achieved with 20% piperadine (a 3/5/3 min cycle performed twice) and coupling through 4 equiv HATU. We analyzed the peptide synthesis at various chain lengths through analytical cleavage. After the coupling of 16AAs, we used pseudo-proline to improve peptide solvation and reduce the aggregation due to the presence of excess hydrophobic AAs (L, I, A, and G). Next, we performed manual deprotection (3/5/3 min) followed by 2.5 h coupling with pseudo-proline of Asn-Ser (Fmoc-Leu-Ser(ψMe,Mepro)-OH, 2.5 equiv) in the presence of 2.5 equiv HATU at position Asn120-Ser121. After two coupling cycles in the synthesiser, Fmoc-Ser(PO3H-Bzl)-OH (2.5 equiv) was manually coupled at position S116 for 2.5 h in the presence of 2.5 and 6 equiv HATU and DIEA, respectively. The next 10 AA couplings (except for Cys109, manual coupling in the presence of HATU for 2.5 h) were performed in the microwave synthesiser (CEM Liberty Microwave Peptide Synthesizer) with the following the settings:

- Solvent volume: Total volume of 4 mL in DMF or NMP
- Coupling: Time 5 min, Power 20W, Temp 75 ºC (Coupling reagent: HATU, 8 equiv)
- Deprotection: Time 3 min, Power 20 W, Temp 75 ºC (20% piperadine in DMF)

The last AA, Boc-Thz, was introduced through manual coupling (1.5 h) in the presence of 2.5 and 6 equiv HATU and DIEA, respectively. The desired peptide with C-terminal thioester was obtained using several steps: (1) the resin was treated with 4-nitrophenylchloroformate (50 mg) dissolved in 3 mL dichloromethane (DCM) for 20 min, and this step was repeated 3 times; (2) the resin was thoroughly washed with DCM followed by DMF; (3) the resin was treated with 10% DIPEA in DMF for 10 min, and this step was repeated 3 times; (4) the resin was treated with a solution (10 mL) of 90% TFA, 5% DCM, 2.5% TIPS, and 2.5% water for complete cleavage of peptide from the resin (3.5 h); and (5) the solution was directly added to cold ether to obtain the precipitate from the crude peptide (60 mg crude). As the crude peptide had limited solubility in the HPLC buffers, any further purification was abandoned.

**6.2 Synthesis of Thz-Htt106-137-Nbz-pT107**. Thz-Htt106-137-Nbz-pS107 was synthesized in a manner similar to that of Thz-Htt106-137-Nbz-pS116 with several changes (Figure 8). After the coupling of pseudo-proline at position Asn120-Ser121, the synthesis was continued (except for Cys109, manual coupling in the presence of HATU for 2.5 h) in the microwave synthesiser. Fmoc-Thr (PO3H-Bzl)-OH (2.5 equiv) was manually coupled for 2.5 h in position

T107 in the presence of 2.5 and 6 equiv of HATU and DIEA, respectively. Finally, after resin cleavage, 45 mg of the crude peptide was obtained. As the crude peptide had limited solubility in the HPLC buffers, no further purification was undertaken.

**7. Dot blot analysis to validate Anti-pT107 and Anti-pS116 antibodies**

The crude mixture resulting from peptide synthesis of Thz-Htt106-137-Nbz-pT107 and Thz-Htt106-137-Nbz-pS116 was used to validate the antibodies. 100 µg of crude peptides were initially dissolved in DMSO followed by the addition of Tris buffer (10 mM Tris, 150 mM NaCl, pH 7.4) to yield a final concentration of 5% DMSO in the solution. Five microliter samples of each peptide were spotted on the nitrocellulose membrane and incubated in Odyssey blocking buffer for 60 min at room temperature. The blocking buffer was poured off, and the primary antibody (Anti-pT107, 1:500 or Anti-pS116, 1:500) was added for 2 h at room temperature in 1 X PBS buffer. The membranes were washed (3 X 10 min) with PBS-Tween 1X (PBST) followed by incubation with secondary goat anti-mouse Alexa680 (1:10000) for 1 h at room temperature. The membranes were washed again three times with PBST, and protein bands were then detected and developed using an Odyssey® CLx System as shown in Figure S9.

**8. Kinase screening for identifying site-specific kinases**

Htt105-171 (210 ug) was dissolved in 218.5 ul of kinase buffer (50 mM Tris, 25 mM $MgCl_2$, 8 mM EGTA, 4 mM EDTA, 2 mM DTT). The reaction mixture was thoroughly vortexed to completely dissolve the peptide. The pH was then adjusted to 7, and 11.5 ul of 100 mM of ATP (5 mM final concentration) was added. Aliquots of 30 ul of the reaction mixture were prepared, and 28 ul of MilliQ water was added to each aliquot. A different kinase (88 kinases were screened) was added to each aliquot and the mixture was incubated at 30 $^{o}$C (Figure S10). The final concentration of the Htt (105-171) was 0.5 ug/ul, and the final volume was 60 ul in each aliquot. The samples were analyzed using ESI-MS and dot blot using Antip-T107 and Anti-pS116 antibodies. To validate the site of phosphorylation by kinases, the screening was repeated with Htt105-171-T107A, Htt105-171-S116A, and Htt105-171-S120A proteins following the same protocol detailed above.

**MS and dot blot analysis for kinases screening** An aliquot of 5 ul of the sample was diluted four times with $H_2O$ (0.1% TFA), and 10 ul were analyzed via ESI-MS. The kinase reaction was monitored by MS over time (t = 0, 4, 6 h and overnight). The samples were also analyzed through dot blot using Anti-pT107 and Anti-pS116, as mentioned previously.

**9. Determining the NLK phosphorylation site on Htt105-171**

A procedure similar to the one previously discussed for kinase screening was followed. Briefly, 50 ug Htt105-171 protein (Htt105-171, Htt105-171-T107A, Htt105-171-S116A, or Htt105-171-S120A) was dissolved in 200 ul of kinase buffer. The pH was adjusted to 7, and 5 ul of 100 mM of ATP was added. NLK was then introduced, and phosphorylation was monitored via ESI-MS at 2, 4, and 6 h (Figure S11).

**10. Semi-synthesis of Htt105-171-pT107, Htt105-171-pS116, and Htt105-171-pT107-pS116**

Htt105-171 (6 mg) was dissolved in buffer containing 5.7 ml, 100 mM Tris, 50 mM $MgCl_2$, 16 mM EGTA, 8 mM EDTA, and 4 mM DTT. The protein was completely solubilized to obtain a clear solution and then adjusted to a pH between 7.0 and 7.1 using 1 M HCl. ATP solution (pH 7) was added followed by TTBK1 kinase (200 ug, 1 ug for 30 ug peptide) for T107, MLK3 (300 ug, 1 ug for 20 ug for peptide) for S116, and MST3 (200 ug, 1 ug for 30 ug for peptide) for T107-S116. The pH of the reaction mixture was checked and adjusted to pH 7 to yield the best result. The samples were incubated at 30 °C for 2 h (TTBK1), 6 h (MLK3), and 3 h (MST3), and the progress of the reaction was monitored using UPLC (C4 column, 5%–95%, acetonitrile, 4 min) and LC/MS. After the completion of the reaction, the crude protein was purified via RP-HPLC using a semi-preparative C4 column (Phenomenex or Higgins, 5 µm, 300Å, 10 x 250 mm). HPLC method used 5% acetonitrile for 10 min, and a gradient of 5%–40% acetonitrile for 2 min, 40%–90% ACN for 50 mins, 90%–95% for 5 min, and 95% ACN for 10 min was used, and Htt105-171-pT107, Htt105-171-pS116, and Htt105-171-pT107-pS116 were eluted at 61, 55, and 54 min, respectively. The eluted fractions were pooled based on their purity as determined by ESI-MS and UPLC analysis. The yields of Htt105-171-pT107, Htt105-171-pS116, and Htt105-171-pT107-pS116 were 2.4, 1.6, and 2 mg, respectively. It was necessary to perform the purification immediately after the reaction to secure high yields.

**11. Semi-synthesis of phosphorylated Htt171-23Q/43Q**

Htt2-104-23Q-Mesna (1.2 equiv) or Htt2-104-43Q-Mesna (2 equiv) was dissolved in TFA (10 ug protein in 1 uL TFA) and incubated for 30 min at room temperature (TFA treatment facilitated disaggregation of the protein and improved solubility of the fragment significantly). Later, under a gentle stream of nitrogen, the TFA was evaporated and the protein was re-dissolved in ligation buffer (8 M Urea, 1 M proline, 50 mM trehalose, 100 mM TCEP, and 50 mM MPAA in MQ water, pH 7.3). The phosphorylated Htt105-171 (1 equiv) was added to the ligation buffer. The pH was adjusted to 7.3, and the reaction was continued at 37 °C. After completion of the reaction, the crude mixture was purified using RP-HPLC with a semi-preparative C4 column (Phenomenex or Higgins, 5 µm, 5 µm, 300Å, 10 x 250 mm). For purification, the HPLC method used 5% ACN for 10 min, 5%–40% acetonitrile for 2 min, 40%–90% acetonitrile for 50 mins, 90%–95% for 5 min, and 95% acetonitrile for 10 min. The eluted fractions were pooled based on their purity as determined by ESI-MS analysis. It was necessary to perform the purification immediately after the ligation reaction to secure high yields (Table 1).

**12. Dot and Western blotting analyses of phosphorylated Htt171-23Q/43Q proteins**

The purified proteins were dissolved in Tris buffer (10 mM Tris, 150 mM NaCl, pH 7.4) to a concentration of 1ug/uL. For dot blot analysis, we followed the procedure mentioned above using primary antibody (Anti-pT107, 1:1000 or Anti-pS116, 1:1000 or Anti-Htt proline-domain, 1:1000) as shown in Figure S15. For western blotting, purified proteins were dissolved in Tris buffer (10 mM Tris, 150 mM NaCl, pH 7.4) to a concentration of 10ug/uL. Ten microliters of each protein sample were loaded on to a 15% polyacrylamide 1.5 mm-thick gel. The protein samples were separated at 150 V for 1 h and then transferred onto nitrocellulose membranes using a semidry transfer system (Bio-Rad). The membrane was blocked in Odyssey blocking buffer for 1 h at room temperature. Membranes were then incubated overnight at 4 °C with primary antibody (Anti-Htt proline-domain, 1:1000). After three washes with PBST buffer, the membrane was incubated with secondary goat anti-mouse Alexa680 (1:10000) for 1 h at room temperature. The membranes were washed again three times with PBST, and proteins were detected and developed using a Li-COR scanner at 700 nm.

**13. *In vitro* aggregation assay**

The proteins were dissolved in HFIP:TFA (1:1, 1 mL for 0.5 mg protein) and incubated for 30 min at room temperature (HFIP and TFA facilitated protein disaggregation and improved solubility). Later, the sample was dried under a gentle nitrogen stream, and the proteins were re-dissolved Tris buffer (10 mM tris) to a concentration of 10 µM. After dissolution, the samples were maintained on ice until the aggregation was initiated. The protein concentration was determined by weight and later adjusted using UPLC with standards of known concentrations. Aggregation was initiated by incubating the proteins at 30 °C. For monitoring the aggregation kinetics of the proteins, 100 µL of 10 µM protein sample was plated in triplicate in a Nunc 96-well fluorescence clear bottom plate. One microliter of ThS (1 mM) was added to all the samples to obtain a concentration of 10 µM in each sample. The ThS fluorescence was then measured using an excitation and emission wavelengths of 440 and 480 nm, respectively, for cycles of 20 min over time without shaking at 30 °C using a Fluostar Omega® plate reader. The raw data were extracted and plotted using GraphPad Prism 8 software.

**14. Circular dichroism spectroscopy measurements**

The purified proteins were dissolved in Tris buffer (10 mM Tris, 150 mM NaCl, pH 7.4) to a concentration of 30 µM (200 uL) and analyzed using a circular dichroism spectrometer. The spectra were recorded in the range of 190 to 280 nm using a 1.0 mm path length quartz cuvette. The data points were acquired in a continuous scanning mode at a speed of 25 nm/min. The processed spectra were obtained by subtracting the baseline signal (quartz cuvette) from the protein spectra with no further smoothening. For thermal denaturation studies, CD for the proteins was measured for heating and cooling cycles. The raw data were converted to mean residue ellipticity ($\theta_{MRW}$) and plotted using Origin 2019 graphing and analysis software. The secondary structure for the phosphorylated Htt105-171 proteins was determined through online CD analysis software (such as Bestsel[1], K2D2[2], and K2D3[3]). The final helical percentage was determined by averaging of the helical percentage obtained from the three software sets.

**15. Sedimentation assay**

The proteins were dissolved in Tris buffer (10 mM Tris, 150 mM NaCl, pH 7.4) to a concentration of 20 µM (500 uL). The concentration of the soluble protein was monitored through UPLC analysis. Aggregation was initiated by incubating the protein at 30 °C. To monitor the soluble fraction, an aliquot (30 µL) of the sample was removed at different time points and centrifuged (4°C, 18000 RCF for 25 min) to remove aggregates. The supernatant was analyzed via UPLC. The area under the curve forming the UPLC peak at t = 0 h was considered 100%, and a change in the area of the peak area was used to calculate the percentage of soluble protein.

**16. Transmission electron microscopy (TEM)**

Five microliters of sample were deposited on a glow discharged (30 sec) 200 mesh formvar-coated TEM grid and incubated for 90 s. The grids were washed three times with 25 µl of Milli-Q water and stained with 3 drops of freshly prepared 2% uranyl acetate for 20 s. The grids were air-dried for 30 min. The samples were imaged using a TEM instrument (FEI Tecnai Spirit) at 80 kV. For time-dependent EM analysis, samples were taken from the aggregating solution at different time points, and EM grids were further prepared as mentioned above.

**17. Sample preparation for Cryo-EM**

A Cu 200 meshes Lacey carbon film grid was glow discharged for 6 seconds and 5 µl of the sample was incubated on the surface for 90 s. The grid was gently washed once with 25 µl of Milli-Q water to not disattach the fibrils and dilute the salt concentration existing in the buffer. The reverse blotting approach was manually performed and the grid rapidly submerged in liquid ethane using a plunge freezer. The grid was then transferred to an FEI Tecnai F20 instrument while it was well-preserved at cryogenic temperature. Imaging was performed at 200 kV from the area with proper ice thickness.

**18. Atomic force microscopy (AFM) imaging**

AFM was performed on mica discs that were positively functionalized with 1% (3-aminopropyl)triethoxysilane (APTS) in aqueous solution for 3 min at room temperature. For time-dependent AFM analysis, 20 µL aliquot of 10 µM Htt171-43Q solution was taken from the aggregating solution at different time points and loaded on the surface of mica discs. Deposition took 3 min and was followed by gentle drying under a nitrogen flow. AFM imaging was performed at room temperature using a Park NX10 operating in true non-contact mode that was equipped with a super sharp tip (SSS-NCHR) Park System cantilever.

### 19. Supporting table and figures

**Table 1**. Ligation time, HPLC elution time, and yield of semisynthetic Htt171-23Q/43Q and phosphorylated Htt171-23Q/43Q proteins.

| Proteins | Ligation time | HPLC elution time | Ligation yield |
|---|---|---|---|
| Htt171-23Q | 4h | 36 ml | 30% |
| Htt171-43Q | 4h | 39 mL | 32% |
| Htt171-23Q-pT107 | 3h | 38 ml | 30% |
| Htt171-23Q-pS116 | 4h | 34 ml | 32% |
| Htt171-23Q-pT107-pS116 | 4h | 35 ml | 34% |
| Htt171-43Q-pT107 | 4h | 42 ml | 24% |
| Htt171-43Q-pS116 | 4h | 39 ml | 25% |
| Htt171-43Q-pT107-pS116 | 4h | 39 ml | 27% |

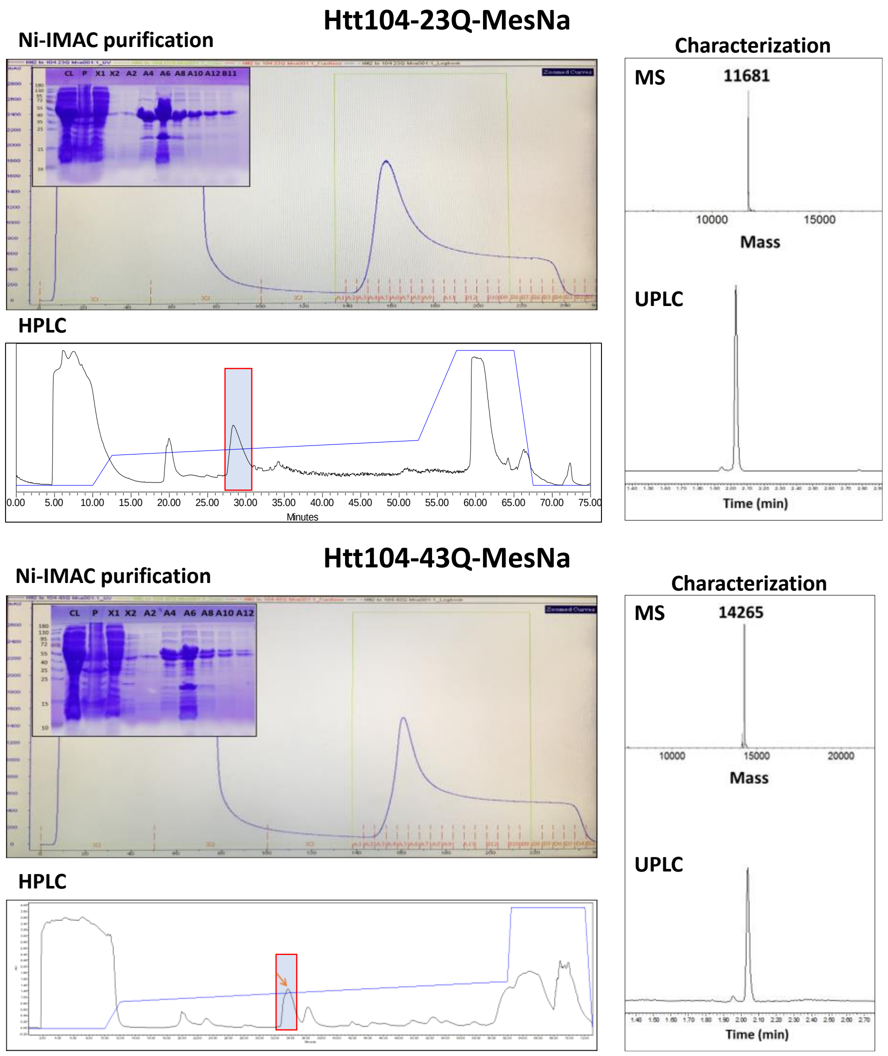

**Figure S1**. Expression and purification of Htt104-23Q-MesNa and Htt104-43Q-MesNa. Representative chromatogram of the Ni-IMAC purification (Inset: analysis by SDS-PAGE of the purification fractions), RP-HPLC chromatogram for the purification and characterization by ESI/MS and UPLC.

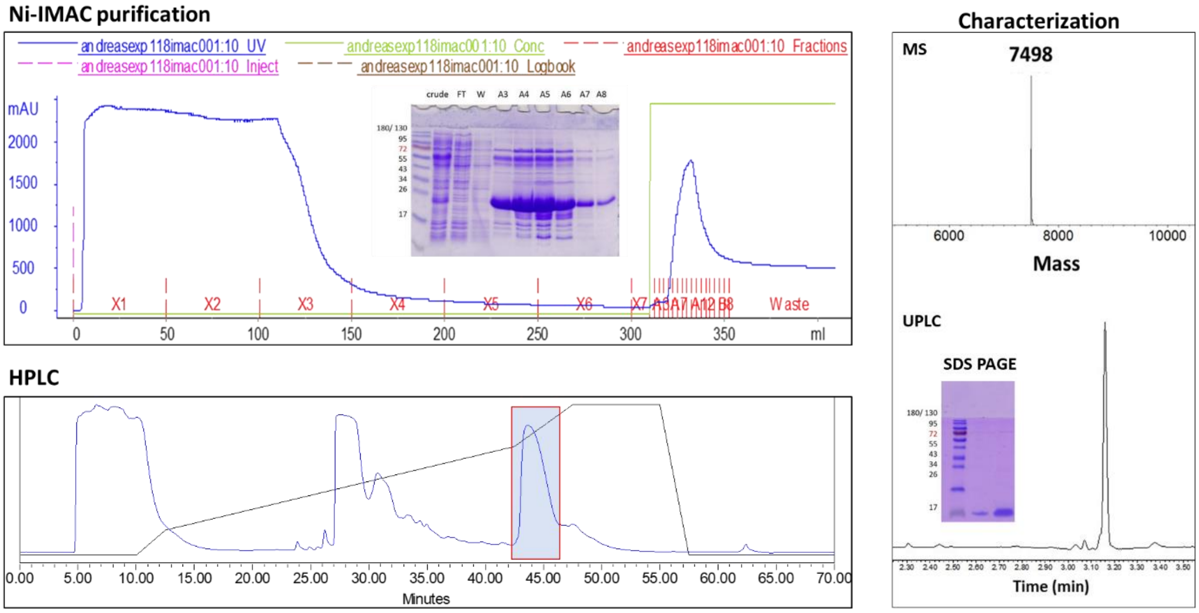

**Figure S2**. Expression and purification of Htt105-171. Representative chromatogram of the Ni-IMAC purification (Inset: analysis by SDS-PAGE of the purification fractions), RP-HPLC chromatogram for the purification and characterization by ESI/MS, UPLC and SDS-PAGE.

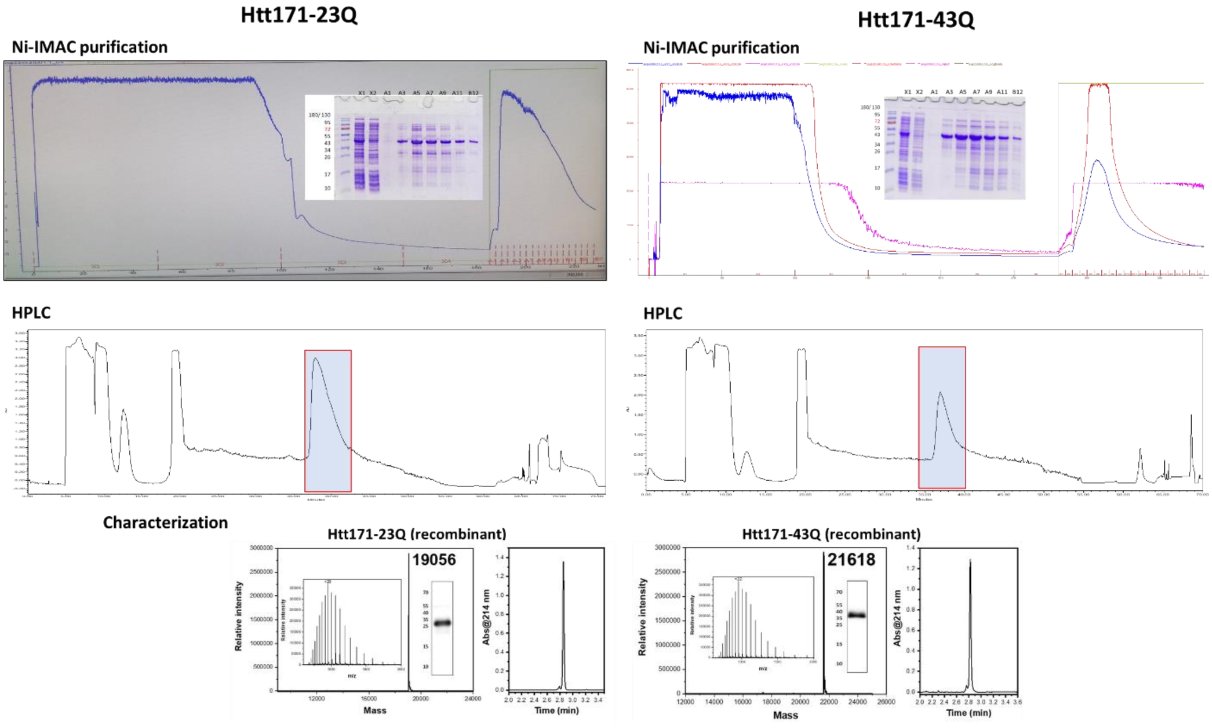

**Figure S3**. Expression and purification of Htt171-23Q and Htt171-43Q. Representative chromatogram of the Ni-IMAC purification (Inset: analysis by SDS-PAGE of the purification fractions), RP-HPLC chromatogram for the purification and characterization by ESI/MS, UPLC and western blot analysis.

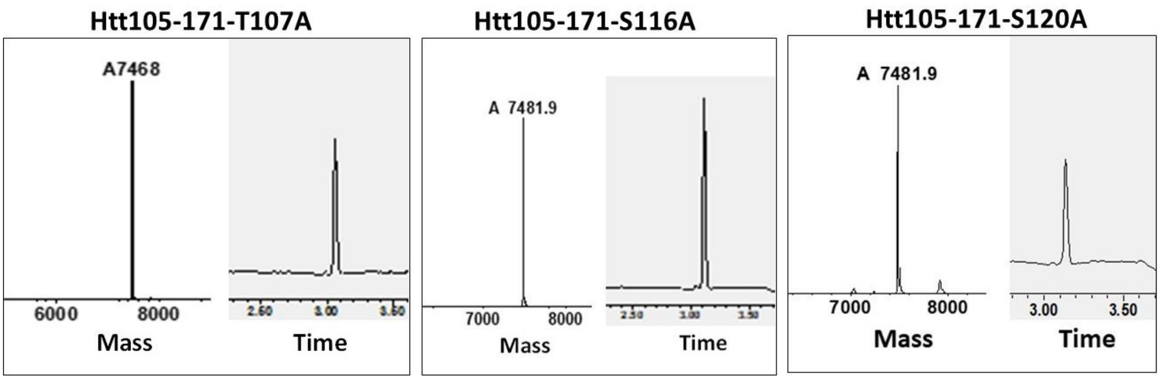

**Figure S4**. Characterization (MS and UPLC) of Htt105-171-T107A, Htt105-171-S116A, and Htt105-171-S120A.

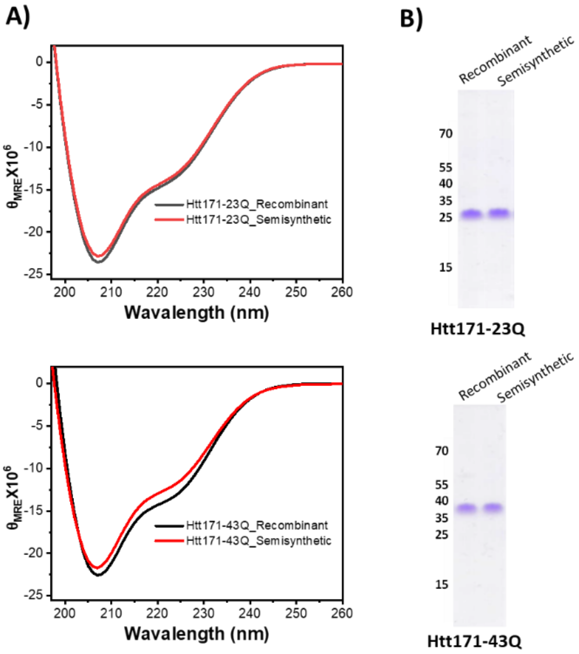

**Figure S5**. Characterization of recombinant and semisynthetic Htt171-23/43Q proteins. **A**) Circular dichroism spectra for the recombinant (black) and semisynthetic (red) Htt171-23Q/43Q. **B**) SDS-PAGE analysis of Htt171-23Q (top) and 43Q (bottom).

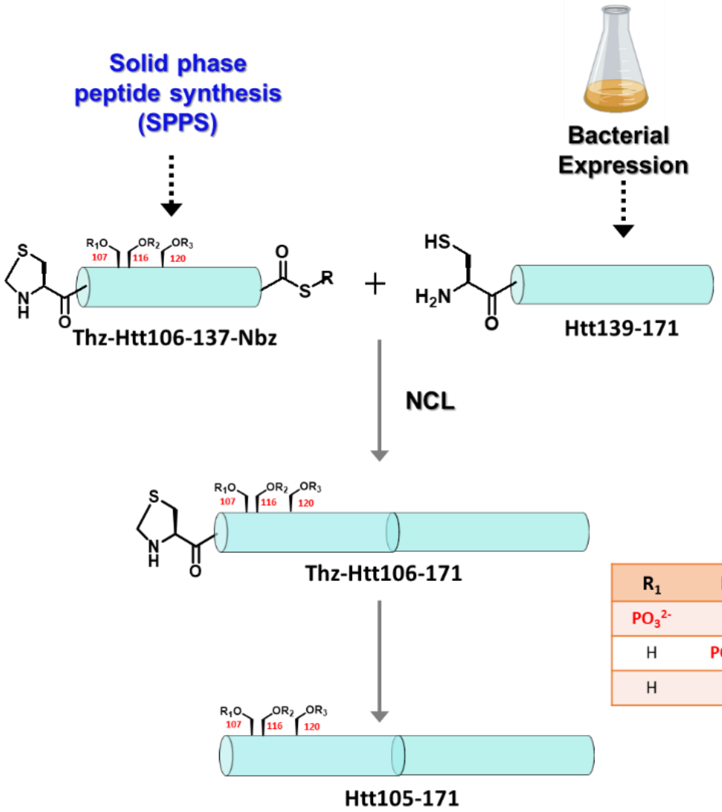

| $R_1$ | $R_2$ | $R_3$ | Protein |
|---|---|---|---|
| $PO_3^{2-}$ | H | H | Htt105-171-pT107 |
| H | $PO_3^{2-}$ | H | Htt105-171-pS116 |
| H | H | $PO_3^{2-}$ | Htt105-171-pS120 |

**Figure S6**. Strategy for the semi-synthesis of phosphorylated Htt105-171 protein through semisynthetic approach.

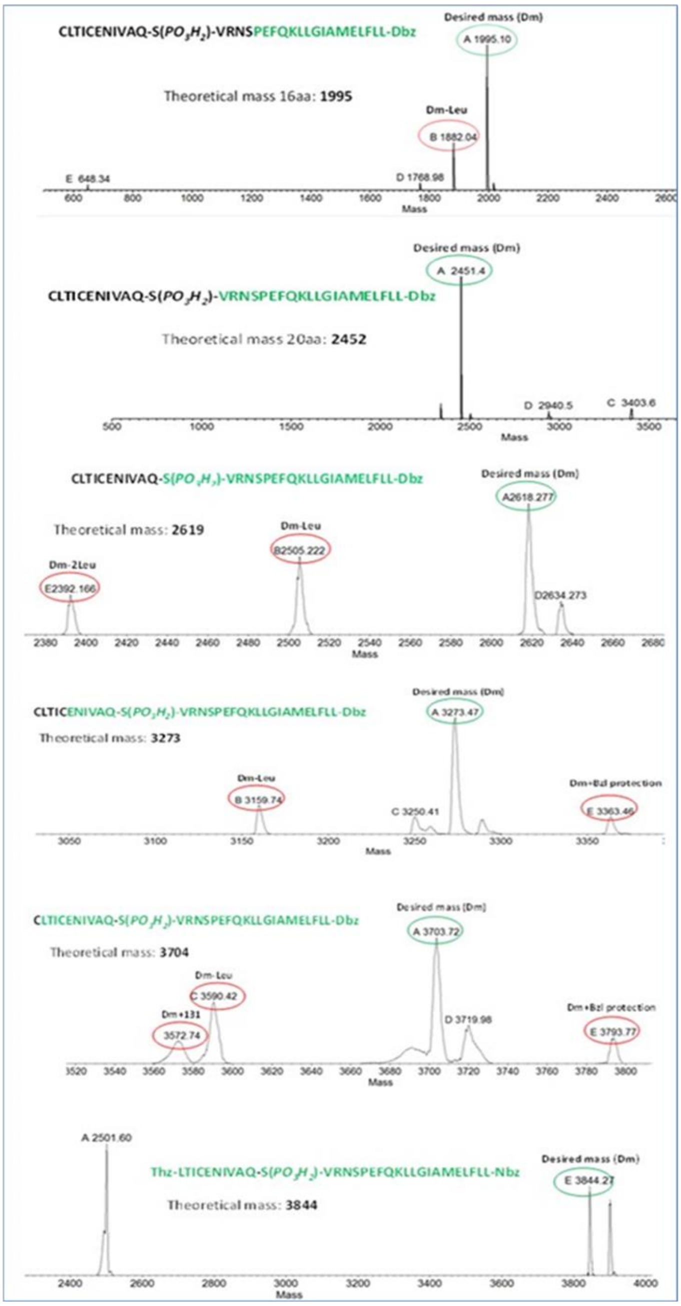

**Figure S7**. SPPS of Thz-Htt106-137-Nbz-pS116 monitored through ESI-MS. Dm= desired mass, Dm-Leu= desired mass with one Leu residue mass less and Dm+Bzl= desired mass + benzyl protection group.

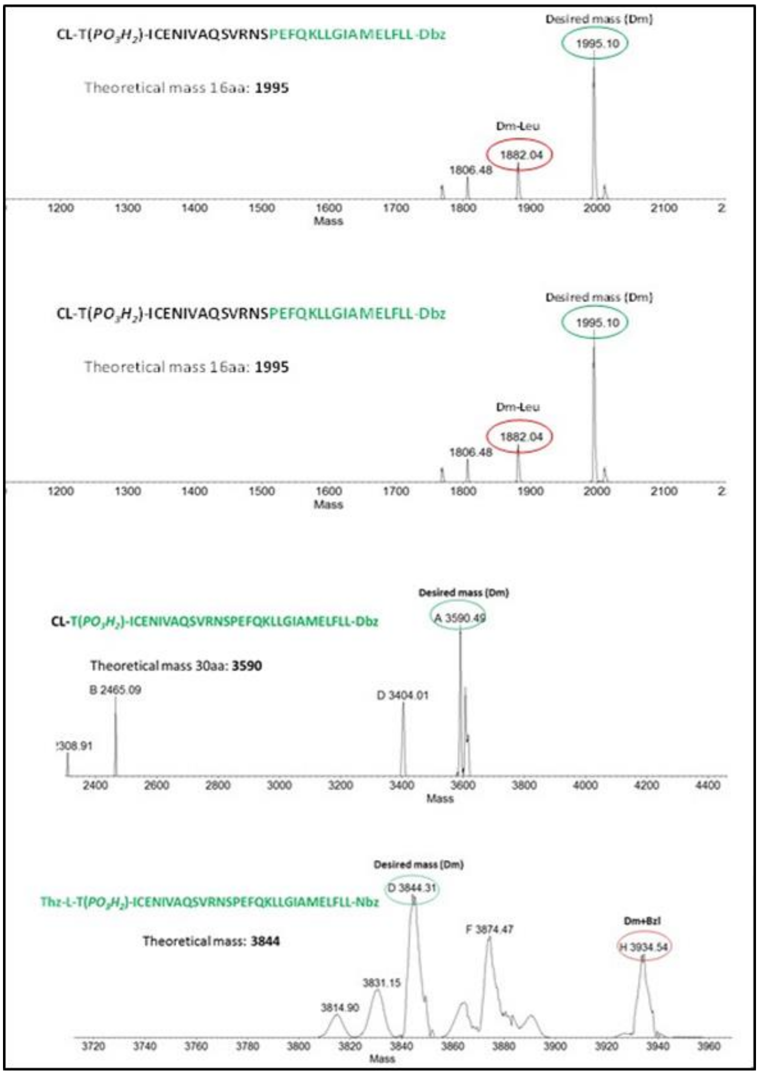

**Figure S8**. SPPS of Thz-Htt106-137-Nbz-pT107 monitored through ESI-MS. Dm= desired mass, Dm-Leu= desired mass with one Leu residue mass less and Dm+Bzl= desired mass + benzyl protection group.

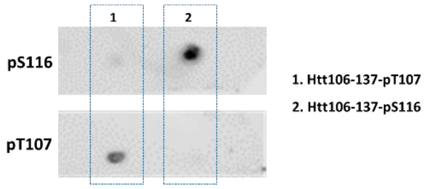

**Figure S9**. Validation of antibodies specific for detecting phosphorylation at T107 and S116 through dot blot analysis.

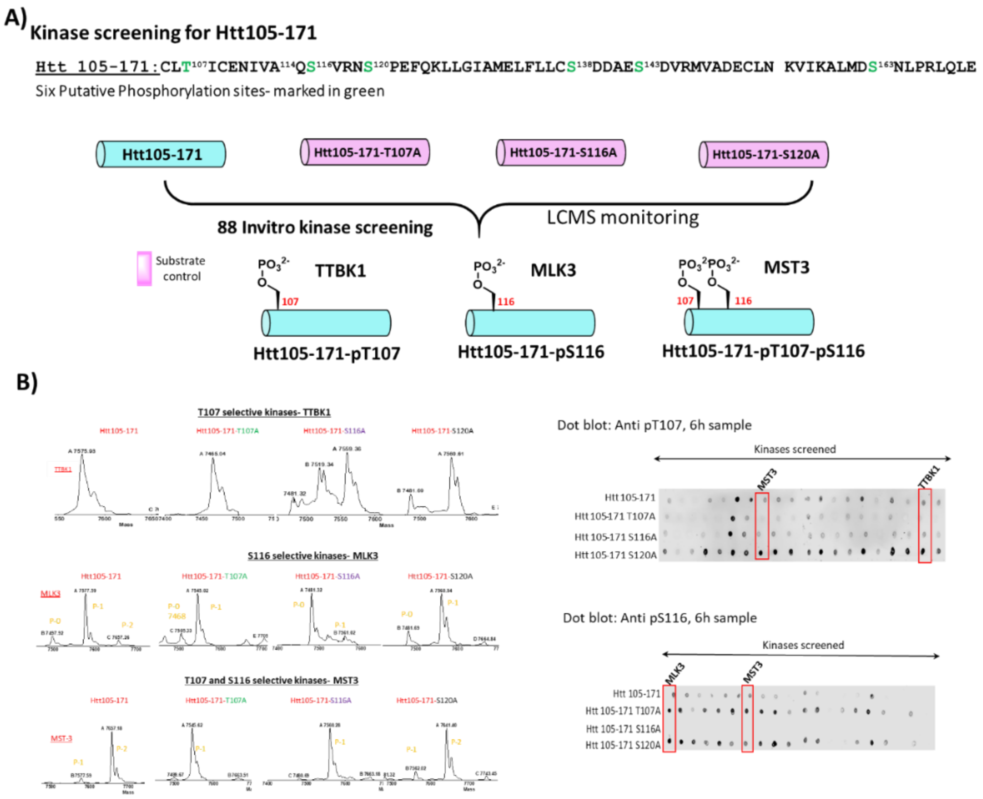

**Figure S10**. Kinase screening for identifying kinases phosphorylating at T107, S116 and S120. A) Outline of kinase screening and the kinases identified for introducing site-specific phosphorylation. B) ESI-MS and Dot blot data for Htt105-171 and mutant Htt105-171 (**4d**, **4e** and **4f**).

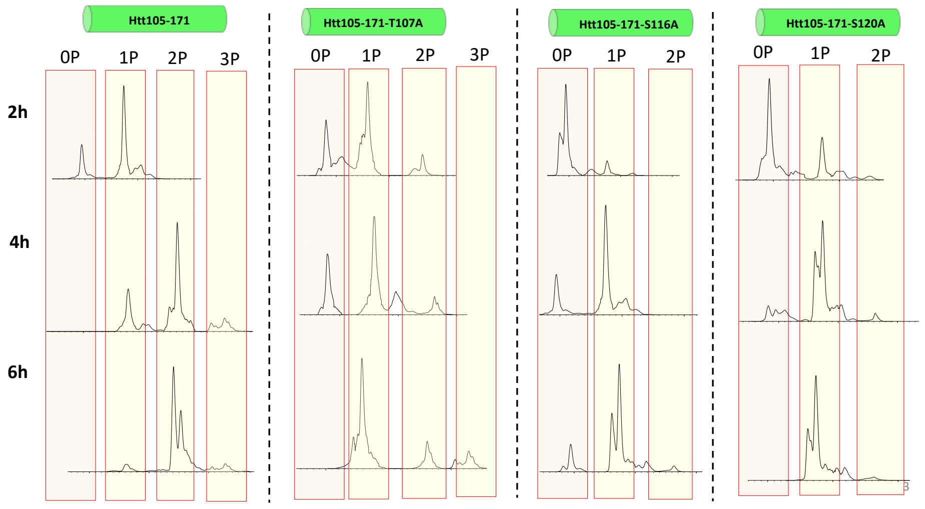

**Figure S11**. Kinase reaction to determine the site of phosphorylation for NLK on Htt105-171 protein. Htt105-171, Htt105-171-T107A, Htt105-171-S116A or Htt105-171-S120A were used in the kinase reaction and phosphorylation was monitored through ESI-MS at 2h, 4h, and 6h.

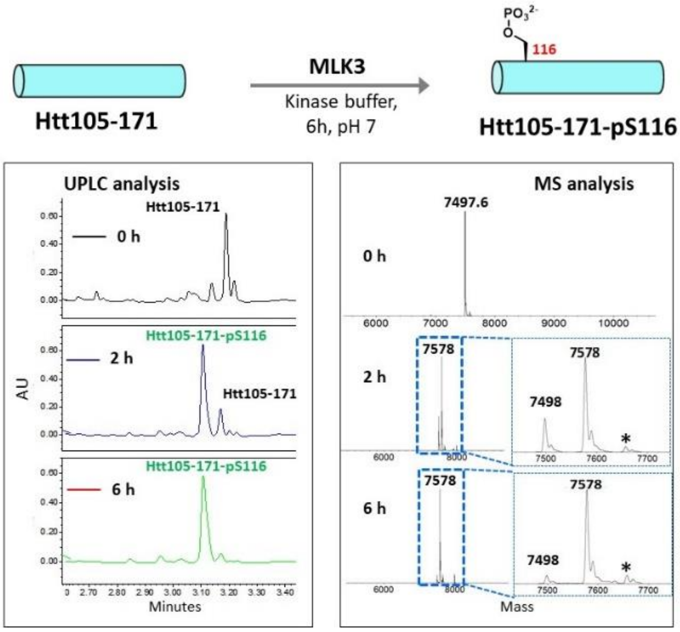

**Figure S12**. Site-specific phosphorylation on Htt105-171 at S116 using MLK3 kinase to obtain Htt105-171-pS116. The reaction was monitored through UPLC (right) and LCMS (left) at 0h, 2h and 6h. * peak corresponds to the diphosphorylated Htt105-171, this impurity was removed during HPLC purification.

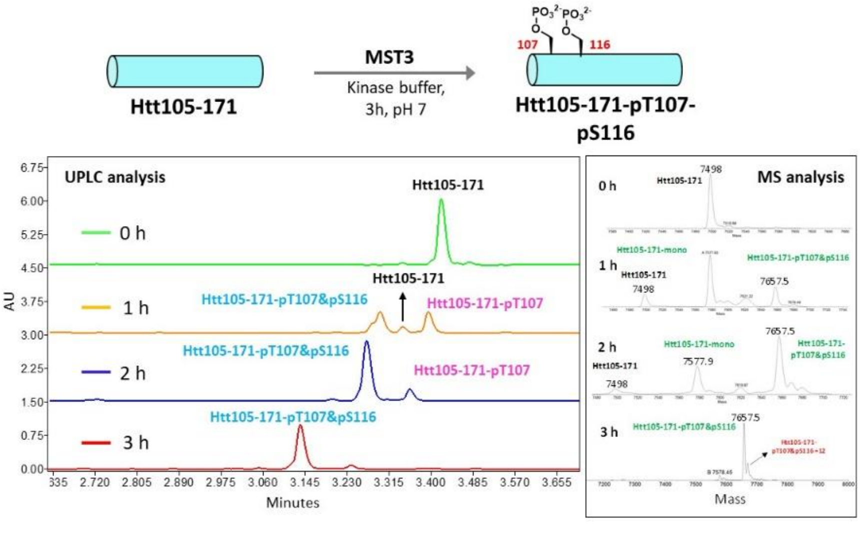

**Figure S13**. Site-specific phosphorylation on Htt105-171 at T107 and S116 using MST3 kinase to obtain Htt105-171-pT107-pS116. The reaction was monitored through UPLC (right) and LCMS (left) at 0h, 1h, 2h and 3h. The highlighted region in UPLC corresponds to the position of Htt105-171 protein.

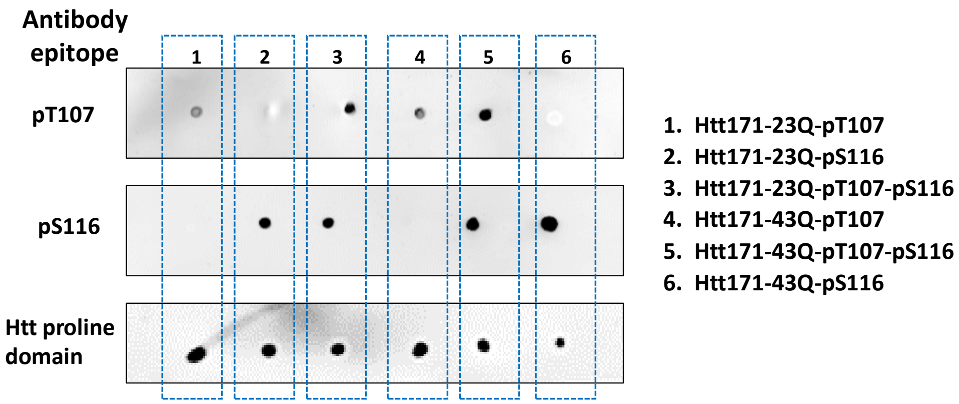

**Figure S14**. Dot blot analysis of semisynthetic phosphorylated Htt171-23Q/43Q proteins to validate the site of phosphorylation on the proteins.

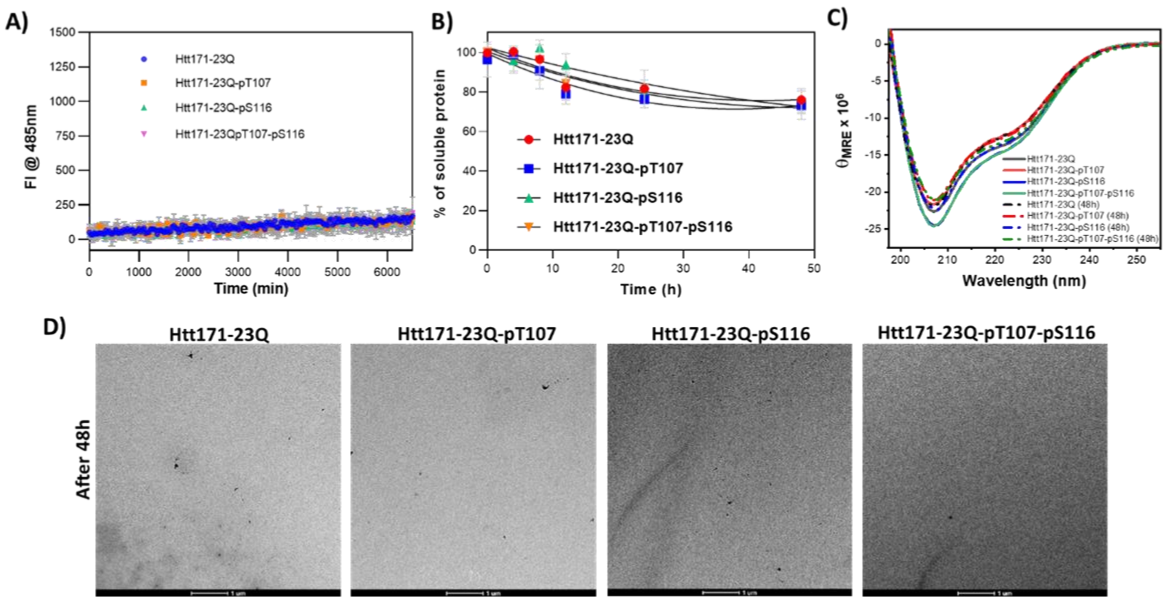

**Figure S15**. *In vitro* aggregation and structure of phosphorylated Htt171-23Q proteins. Aggregation kinetics measured by (A) ThS fluorescence at 485 nm and (B) UPLC sedimentation assay (loss of soluble protein) for the phosphorylated Htt171-23Q (mean ± SEM, n=3). C) CD analysis of phosphorylated Htt171-23Q proteins at 0 and 48h. D) TEM images of the phosphorylated Htt171-23Q proteins at 1 and 48h (scale bars are 500 nm).

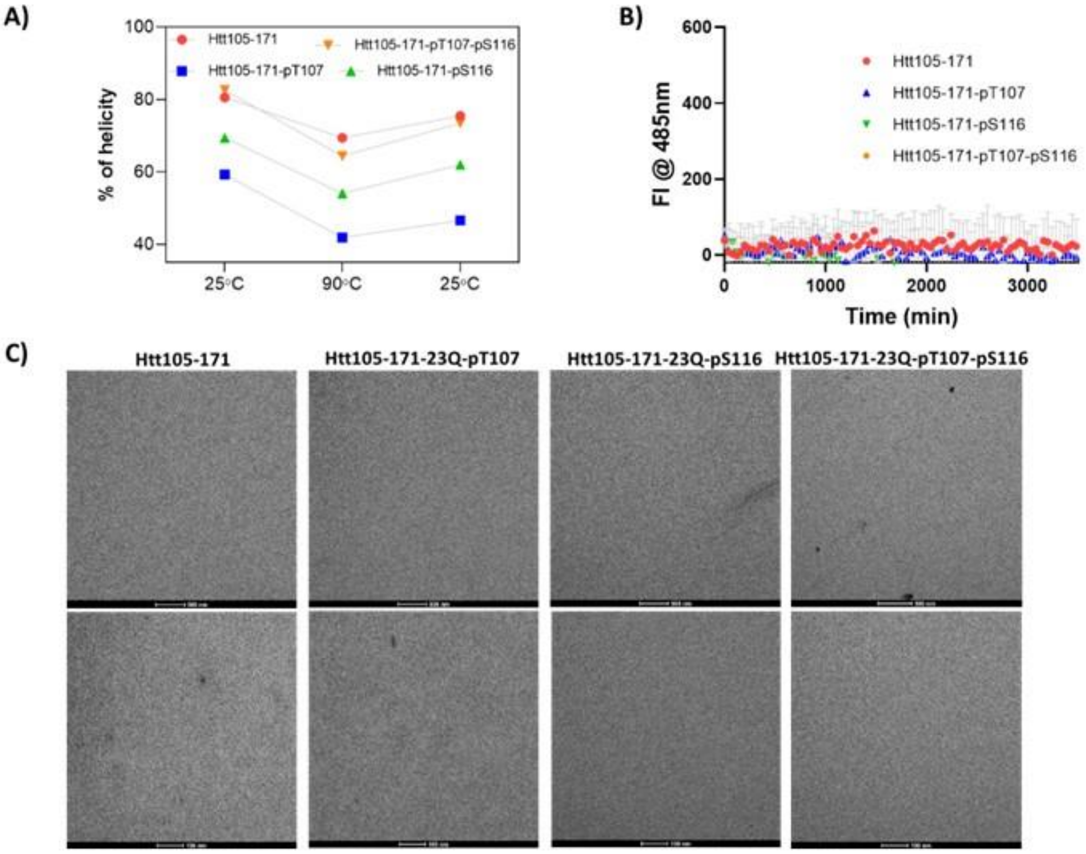

**Figure S16**. A) Percentage of helical conformation of unmodified and phosphorylated Htt105-171 proteins evaluated during thermal denaturation studies. B) Aggregation kinetics measured by ThS fluorescence at 485 nm (mean ± SEM, n=3) for Htt105-171 proteins. C) TEM images of unmodified and phosphorylated Htt105-171 proteins after 48 h (scale bars are 500 nm for the top panel and 100 nm for the bottom panel).

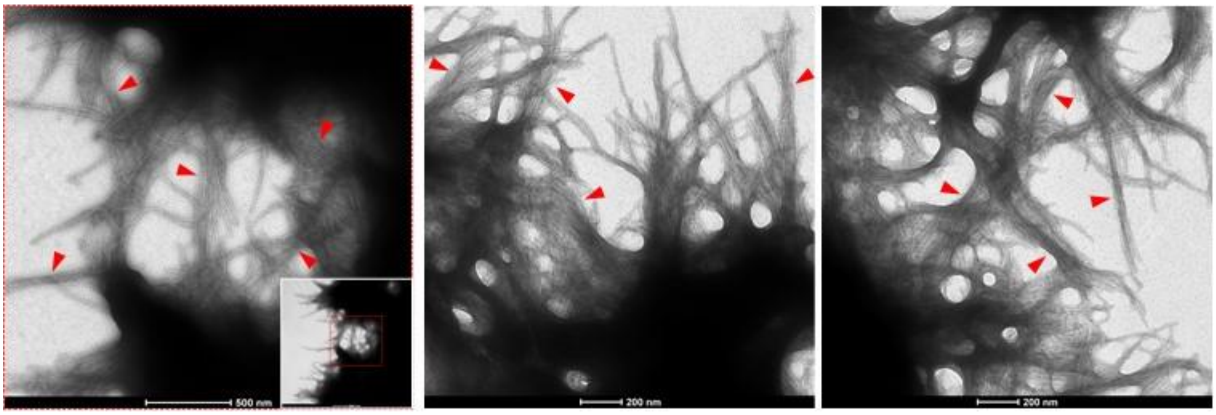

**Figure S17**. EM images showing long fibrils growing on the surface of globular species. Red arrowheads indicate fibrils with high lateral association.
